# *Clostridioides difficile* LuxS mediates inter-bacterial interactions within biofilms

**DOI:** 10.1101/494245

**Authors:** Ross Slater, Lucy Frost, Sian Jossi, Andrew Millard, Meera Unnikrishnan

## Abstract

The anaerobic gut pathogen, *Clostridioides difficile*, forms adherent biofilms that may play an important role in recurrent *C. difficile* infections. The mechanisms underlying *C. difficile* community formation and inter-bacterial interactions are nevertheless poorly understood. *C. difficile* produces AI-2, a quorum sensing molecule that modulates biofilm formation across many bacterial species. We found that a strain defective in LuxS, the enzyme that mediates AI-2 production, is defective in biofilm development *in vitro*. Transcriptomic analyses of biofilms formed by wild type (WT) and *luxS* mutant (*luxS*) strains revealed a downregulation of prophage loci in the *luxS* mutant biofilms compared to the WT. Detection of phages and eDNA within biofilms may suggest that DNA release by phage-mediated cell lysis contributes to *C. difficile* biofilm formation. In order to understand if LuxS mediates *C. difficile* crosstalk with other gut species, *C. difficile* interactions with a common gut bacterium, *Bacteroides fragilis*, were studied. We demonstrate that *C. difficile* growth is significantly reduced when co-cultured with *B. fragilis* in mixed biofilms. Interestingly, the absence of *C. difficile* LuxS alleviates the *B. fragilis* mediated growth inhibition. Dual species RNA-sequencing analyses from single and mixed biofilms revealed differential modulation of distinct metabolic pathways for *C. difficile* WT, *luxS* and *B. fragilis* upon co-culture, indicating that AI-2 may be involved in induction of selective metabolic responses in *B. fragilis*. Overall, our data suggest that *C. difficile* LuxS/AI-2 utilises different mechanisms to mediate formation of single and mixed species communities.

## Introduction

*Clostridiodes difficile (Clostridium difficile)*, an anaerobic, opportunistic pathogen, is the causative agent of *C. difficile* infection (CDI), a debilitating condition with symptoms ranging from mild diarrhoea to severe pseudomembranous colitis. ~453,000 cases of CDI were reported in the United States in 2011 [1] and there have been increasing reports of CDI from different parts of the world [2-4]. Treatment of CDI is complicated by the fact that 20-36% of cases experience recurrence, relapsing after completion of initial treatment [5]. CDI is primarily a hospital-acquired infection with the elderly being at highest risk [6] and has been associated with the disruption of the gut microbiota as a result of the use of broad-spectrum antibiotics. However, more recently, there has been a reported increase in community-acquired cases where patients do not have the typical risk factors such as antibiotic exposure or recent hospitalisation [7].

Colonisation of *C. difficile* and development of CDI is influenced by composition of the native gut microbiota. Broadly, *Bacteroides*, *Prevotella*, *Difidobacterium*, *Enterococcaceae* and *Leuconostocaceae spp*. correlate negatively [8-10], and Lactobacilli, *Aerococcaceae, Enterobacteriaceae*, and *Clostridium* correlate positively to *C. difficile* colonisation and disease [8, 10-13]. While mechanisms underlying colonisation resistance are not entirely clear, some pathways have been described recently. Secondary bile acids produced by bacteria like *Clostridium scindens* can inhibit *C. difficile* growth, while other bile acids such as chenodeoxycholate can inhibit spore germination [14-16]. Studies have shown that the ability of *C. difficile* to utilise metabolites produced by the gut microbiota or mucosal sugars such as sialic acid promote *C. difficile* expansion in the gut [17, 18]. However, gaps still remain in the understanding *C. difficile* interactions with members of the gut microbiota.

Research into CDI has primarily focused on the action of two large toxins [19, 20] that cause tissue damage, neutrophil recruitment and a severe inflammatory response [21]. More recently, a number of factors have been shown to influence adhesion of *C. difficile* to host cells and early colonisation, including cell wall proteins, adhesins and flagella [22-26]. *C. difficile* also produces biofilms that confer increased resistance to antibiotics [27-29] and have recently shown to be associated with *C. difficile* infection *in vivo*, in close association with other commensal gut species [30, 31].

Formation of adherent communities within the gut requires communication between bacteria. For many species, quorum sensing (QS) is important for the construction and/or dispersal of biofilm communities [32], with bacteria utilising diverse QS systems [32, 33]. Many bacteria possess the metabolic enzyme LuxS, which is involved in the detoxification of S-adenoslylhomocysteine during the activated methyl cycle. Whilst catalysing the reaction of S-ribosylhomocysteine to homocysteine, LuxS produces the bi-product 4,5-dihydroxy-2,3-pentanedione (DPD). DPD is an unstable compound that spontaneously cyclises into several different forms. These ligands are collectively known as autoinducer-2 (AI-2), a group of potent, crossspecies QS signalling molecules [34]. In many bacteria, including *C. difficile*, AI-2 plays a role in biofilm formation, with *luxS* mutants showing a defect during biofilm formation and development [27, 35-40]. The precise mode of action for LuxS in *C. difficile* has remained elusive as a result of conflicting studies and the lack of a clear receptor for AI-2 [35, 41, 42].

Here we investigate the role of LuxS within *C. difficile* and mixed biofilm communities. Interestingly, we find that *C. difficile* LuxS/AI-2 mediates the induction of two putative *C. difficile* R20291 prophages within *C. difficile* biofilms. In mixed biofilms, we show that in the presence of *B. fragilis*, a gut bacterium, *C. difficile* growth is inhibited and this inhibition is alleviated in the absence of LuxS. Dual species transcriptomics show that distinct metabolic pathways are triggered in mixed cultures with the wild type (WT) and *luxS* mutant *C. difficile* strains.

## Results

### LuxS mediates biofilm formation *in vitro*

We previously reported that a R20291 *C. difficile luxS* mutant (*luxS*) was defective in biofilm formation as measured by crystal violet (CV) staining [27]. However, in our subsequent studies, although we see a reduction in biofilms at 24 h, we observe a high variability in the WT biofilms formed at 24 h and in the reduction in the *luxS* mutant between experiments (Figure 1A). Nevertheless, the biofilm defect for *luxS* was very consistent at later time points (72 h) (Figure 1A, B). In spite of differences in total biofilm content, colony counts from the WT and *luxS* biofilms were similar at both times (Figure 1C).

**Figure 1.**
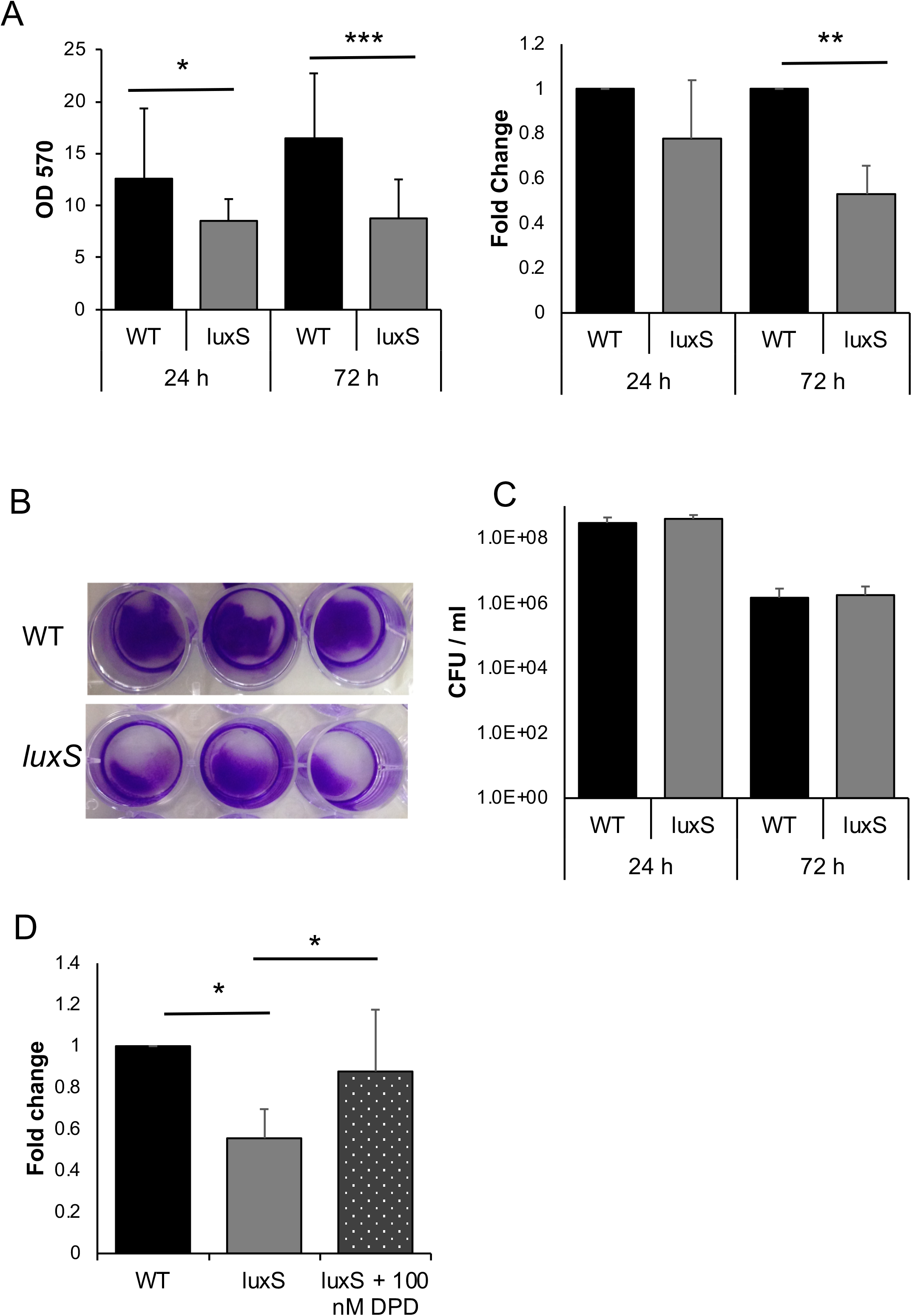
*LuxS* biofilm defect is reversed by addition of DPD. (**A**) WT and *LuxS* biofilms were grown for 24 h or 72 h and stained with 0.2% CV, followed by measuring OD_570_, N=5. (**B**) Colony counts from biofilms (N=7) after 24 h and 72 h. (**C**) Representative pictures of crystal violet stained *C. difficile* WT and *luxS* biofilms after 72 h. (**D**) The AI-2 precursor, DPD, was exogenously supplemented to *LuxS* at a concentration of 100 nM, followed by biofilm staining and quantitation with 0.2% CV after 72 h. Error bars indicate SD, *p < 0.05, ***p<0.001 as determined by students t-test or by Mann-Whitney U test.

To determine if AI-2 signalling is involved in biofilm formation, we first performed an AI-2 assay from both planktonic and biofilm supernatants as described by Carter *et al*. 2005 (Figure S1A). AI-2 is produced maximally in mid-log and stationary phases as previously reported [44]. The WT strain produced less AI-2 in 24 h biofilms compared to log phase culture, while the *luxS* strain did not produce AI-2 as expected (Figure S1B). To study if the biofilm defect could be complemented by chemically synthesised 4,5-dihydroxy-2, 3-pentanedione (DPD), the precursor of AI-2, was supplemented in the culture medium. Whilst high concentrations (>1000 nM) appeared to have no effect on biofilm formation (data not shown), a concentration of 100 nM was capable of restoring the WT phenotype (Figure 1D), indicating that AI-2 may be involved in signalling within *C. difficile* biofilms.

### RNA-seq analysis reveals LuxS-mediated prophage induction

To investigate mechanism of action of LuxS/AI-2 in *C. difficile*, transcriptional profiles of planktonically cultured *C. difficile* WT and *luxS* strains were first compared using RNA-seq. However, surprisingly, no differential transcriptional changes were observed (Accession number E-MTAB-7486). Following this, an RNA-seq analysis was performed with total RNA isolated from *C. difficile* WT and *luxS* biofilms grown in BHIS + 0.1 M glucose for 18 h (Accession number E-MTAB-7523).

The DESeq2 variance analysis package [45] was used to identify genes that were differentially expressed in *luxS* ≥ 1.6-fold relative to the WT strain, with a *p*-adjusted value ≤ 0.05. This pairwise analysis identified 21 differentially expressed genes (Figure 2A & C) (Table 1). Interestingly, all 18 down-regulated genes correspond to two prophage regions located within the *C. difficile* R20291 genome (Figure 2B), as identified using the online phage search tool, Phaster [46, 47]. A Fisher’s exact test (*p*-value < 0.001 for both prophage regions) further confirmed an enrichment of differently regulated genes in prophage regions. There were only three genes upregulated in the *luxS* biofilms compared to the WT; two of these were involved in trehalose utilisation.

**Figure 2.**
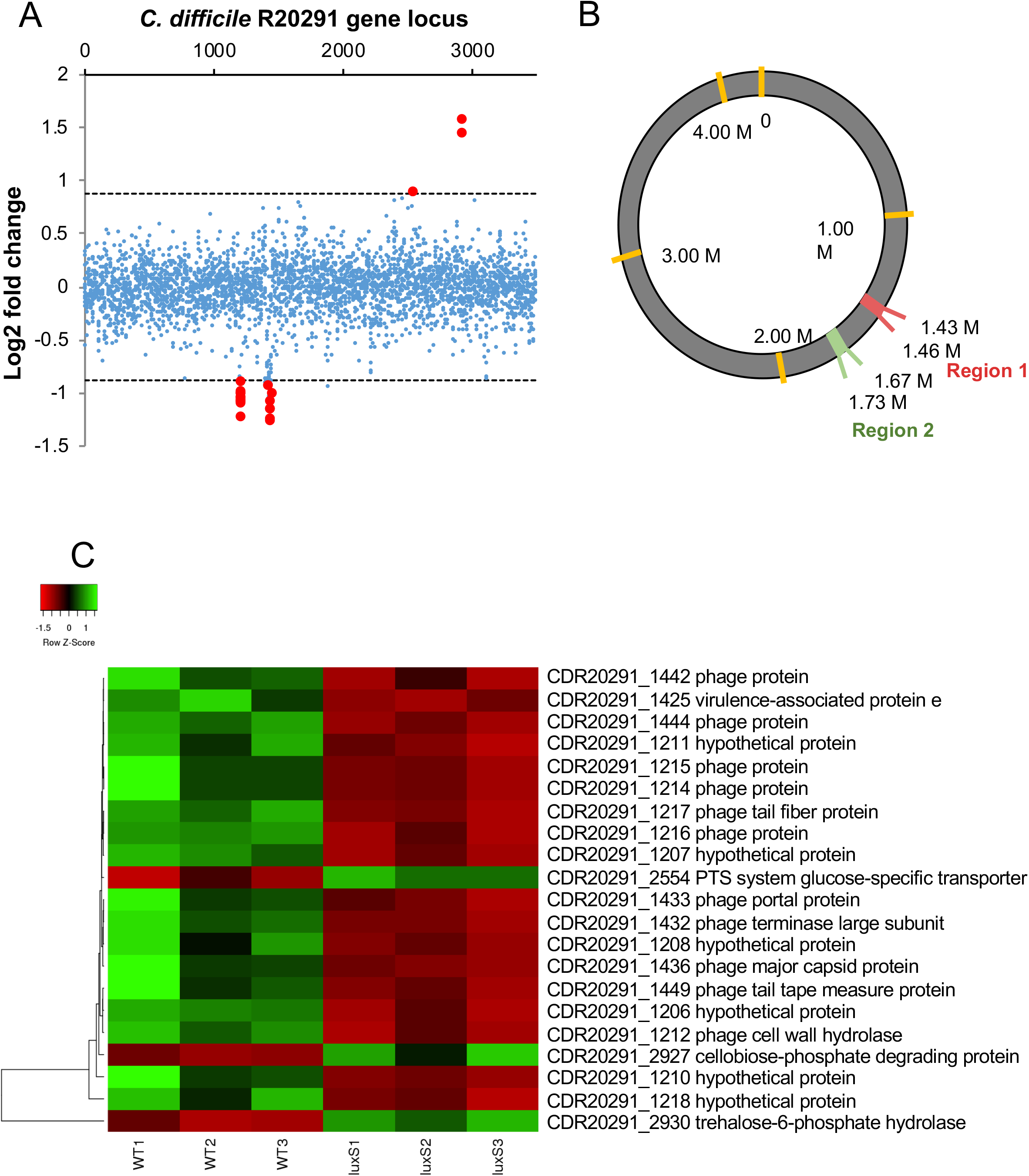
Down-regulation of prophage genes in *C. difficile* luxS mutant. (**A**) Pairwise analysis identified 21 differentially expressed genes in *luxS* (red points). All 18 down-regulated genes clustered into two regions. (**B**) Three prophage regions are identified in the *C. difficile* genome using Phaster. Regions 2 and 3 were down regulated in *luxS*. (**C**) Heat map representation of the genes that were differentially expressed in *luxS* red and green indicate down-and up-regulation respectively when compared to WT. 18 prophage genes were found to be down-regulated in *luxS* relative to WT, whilst three metabolic genes involved in trehalose metabolism were up-regulated in *luxS* relative to WT. Data shown is the mean of 3 independent experiments in triplicates. Differential expression was defined as ≥ 1.6-fold change relative to WT with an adjusted *p*-value ≤ 0.05.

**Table 1.** Genes up- and down-regulated in ***luxS*** relative to the WT *C. difficile*

| No | Gene ID | log2 FoldChange | Gene annotation |
| --- | --- | --- | --- |
| 1 | CDR20291_1206 | -0.994213972 | hypothetical protein |
| 2 | CDR20291_1207 | -1.059645046 | hypothetical protein |
| 3 | CDR20291_1208 | -1.227114763 | hypothetical protein |
| 4 | CDR20291_1210 | -1.08584066 | hypothetical protein |
| 5 | CDR20291_1211 | -1.0763802 | hypothetical protein |
| 6 | CDR20291_1212 | -1.009946545 | phage cell wall hydrolase |
| 7 | CDR20291_1214 | -1.09972952 | phage protein |
| 8 | CDR20291_1215 | -1.038498251 | phage protein |
| 9 | CDR20291_1216 | -1.047731727 | phage protein |
| 10 | CDR20291_1217 | -0.999294901 | phage tail fiber protein |
| 11 | CDR20291_1218 | -0.898724949 | hypothetical protein |
| 12 | CDR20291_1425 | -0.928686375 | virulence-associated protein e |
| 13 | CDR20291_1432 | -1.243533992 | phage terminase large subunit |
| 14 | CDR20291_1433 | -1.154529376 | phage portal protein |
| 15 | CDR20291_1436 | -1.156306053 | phage major capsid protein |
| 16 | CDR20291_1442 | -1.085573967 | phage protein |
| 17 | CDR20291_1444 | -1.257181603 | phage protein |
| 18 | CDR20291_1449 | -1.008627768 | phage tail tape measure protein |
| 19 | CDR20291_2554 | 0.895993497 | PTS system glucose-specific transporter subunit IIA |
| 20 | CDR20291_2927 | 1.587075038 | cellobiose-phosphate degrading protein |
| 21 | CDR20291_2930 | 1.444846424 | trehalose-6-phosphate hydrolase |

Note: P-adjusted value $\leq 0.05$

To demonstrate the presence of phage in the biofilm, cell-free supernatants were treated with DNase, before a subsequent DNA extraction was performed. As the bacterial cells were already removed, only DNA within intact bacteriophages would be protected from DNase. A 16S rRNA gene PCR was performed to confirm digestion of all free extracellular genomic DNA from the biofilm (Figure 3A). PCRs with primers corresponding to genes specific to each prophage, confirmed that the DNA extracted had come from the phage (Figure 3 B). Since cell lysis is linked to phage release, we quantified and compared the total extracellular DNA (eDNA) content of *luxS* mutant and WT biofilms. The WT biofilms contained more eDNA compared to the *luxS* mutant (Figure 3C). Overall, these data suggest that AI-2 may play a role in inducing prophages in *C. difficile* biofilms, which leads to phage-mediated host cell lysis and may contribute to subsequent biofilm accumulation.

**Figure 3.**
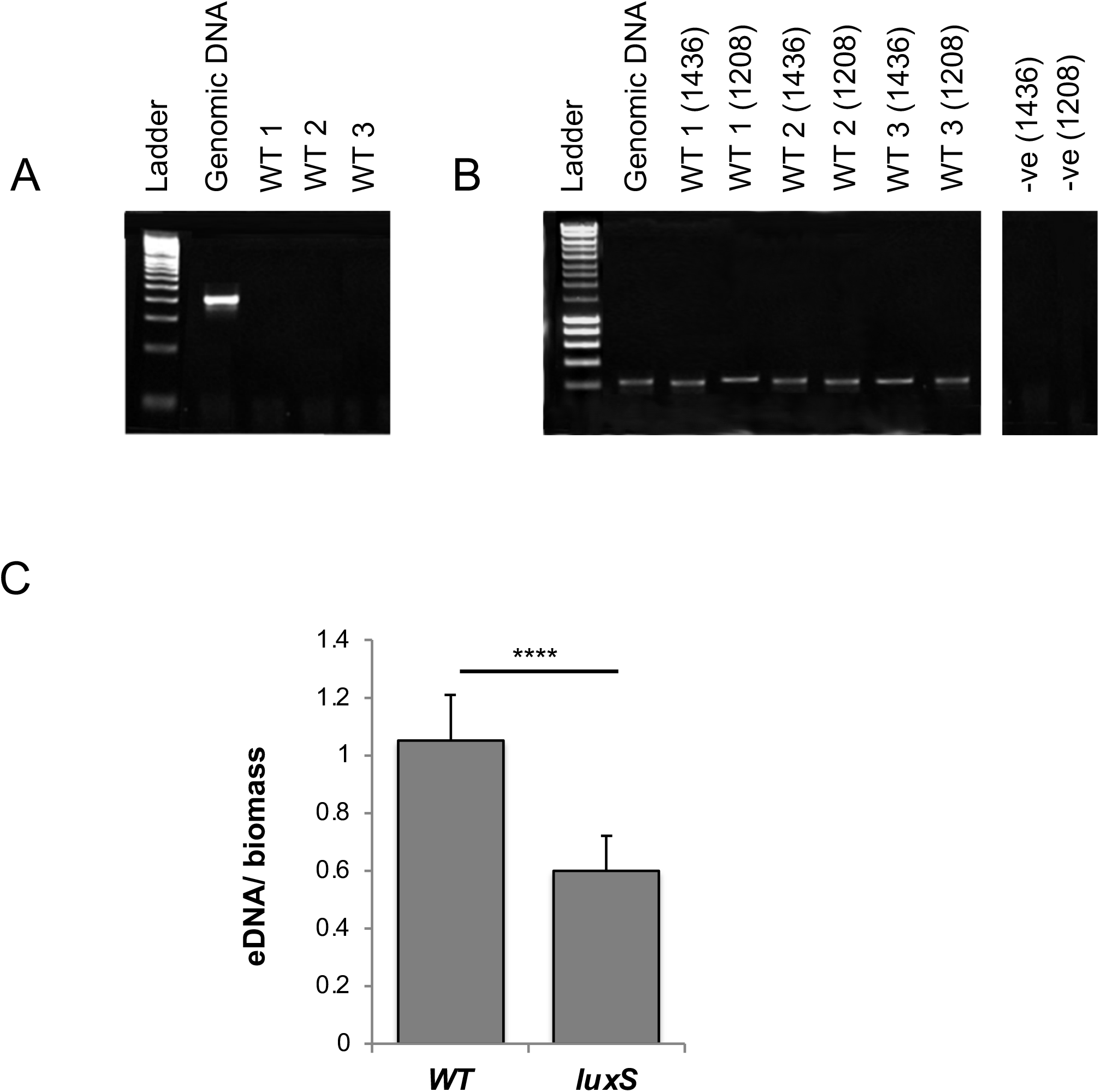
Presence of phage and eDNA in *C. difficile* biofilms. The phage origin of DNA isolated from WT biofilms was confirmed by PCR, using primers for 16S (**A**) and two phage genes (CDR20291_1436 and CDR20291_1208 (**B**). WT-1-3 are three biological replicates. (**C**) Total eDNA extracted from the WT and *luxS* mutant biofilms after 24 h, normalised to the biofilm biomass. N=3, ****p<0.0001 as determined by Mann-Whitney U test.

### *C. difficile* is inhibited when cultured with *B. fragilis* in mixed biofilms

Given the high microbial density within the gut, *C. difficile* likely needs to interact with other members of the gut microbiota to establish itself within this niche. As an interspecies signalling function has been previously proposed for AI-2 [48, 49], we sought to investigate the interactions between a gut-associated *Bacteroides spp* and *C. difficile*.

C. *difficile* formed significantly more biofilms *in vitro* compared with *Bacteroides fragilis* in monocultures, as measured by CV staining (Figure 4A). When both organisms were co-cultured, less biofilm was formed compared to *C. difficile* monoculture (Figure 4A). To investigate the impact of co-culturing on both *C. difficile* and *B. fragilis*, bacterial numbers (CFU/ml) were determined from monoculture and mixed biofilms (Figure 4B). Colony counts obtained from the mono and co-culture biofilms, confirmed that *B. fragilis* was a poor biofilm producer when cultured alone. Interestingly, when both species were co-cultured, the CFU/ml for *C. difficile* was significantly reduced, and the CFU/ml of *B. fragilis* significantly higher. This reduction of colony counts of *C. difficile* was observed at both 24 h (Figure 4B) and 72 h (Figure S2). AI-2 production from single and mixed biofilms was also quantitated. We observed no production of AI-2 by *B. fragilis*, and a reduction of AI-2 production by the mixed biofilms compared with the WT *C. difficile* biofilms (Figure S3). Confocal microscopy analysis of mono and co-culture biofilms showed that the single species biofilms contained more live bacteria compared to co-culture biofilms, which had higher numbers of dead cells (Figure 4C). These data suggest that the presence of *B. fragilis* in biofilms results in inhibition of *C. difficile* growth.

**Figure 4.**
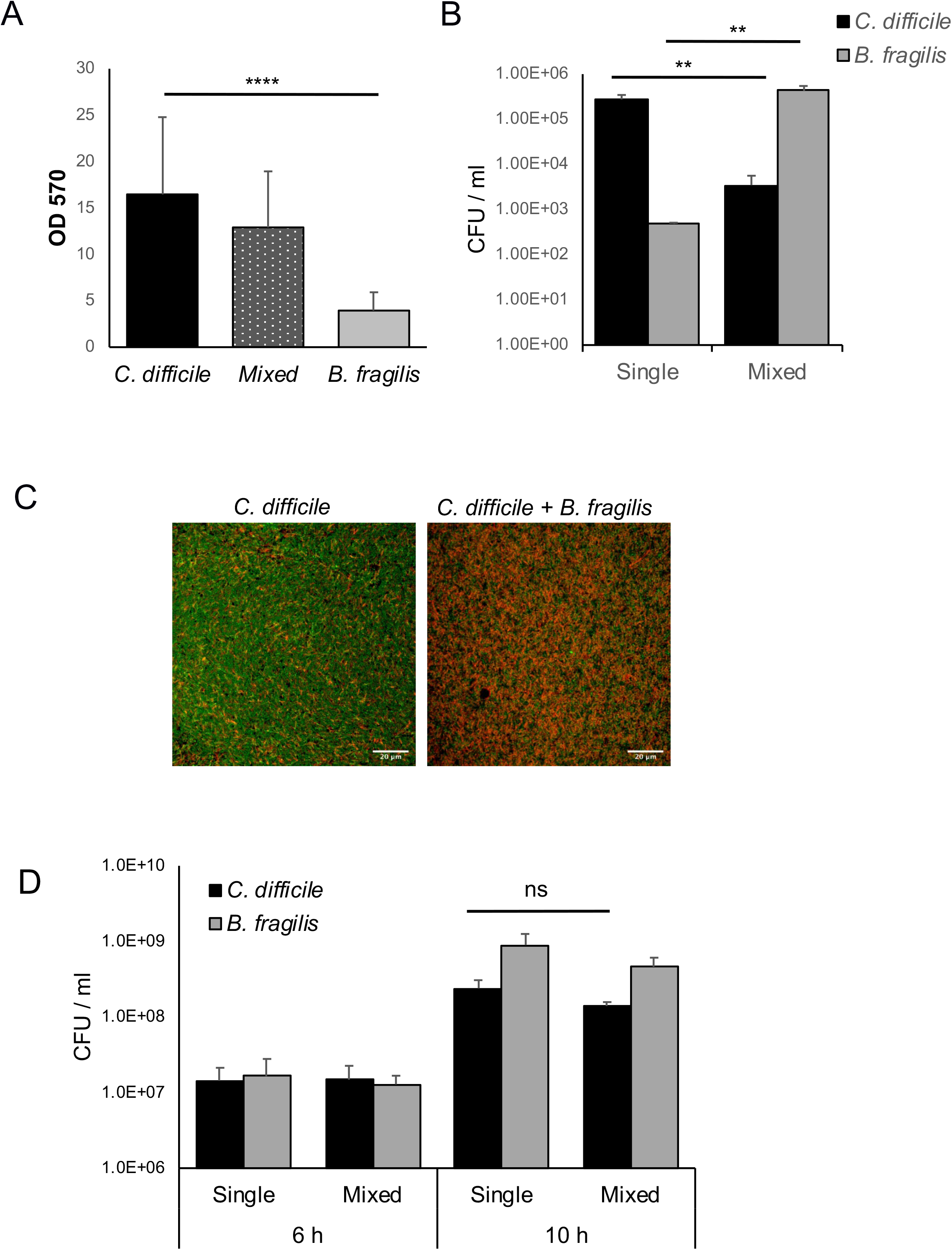
*B. fragilis* mediated inhibition of *C. difficile* in mixed biofilms. (**A**) Biofilm of *C. difficile*, *B. fragilis* and both species co-cultured (mixed) were grown for 24 h and stained with 0.2% CV, followed by measuring OD_570_. (**B**) Colony counts for both *C. difficile* and *B. fragilis* from mono and co-culture biofilms after 24 h. (**C**) Mono and co-culture biofilms were visualised by confocal microscopy after 24 h culture using a live/dead stain. Live bacteria are stained green and dead bacteria stained red using Syto 9 and propidium iodide dye respectively. (**D**) Colony counts for both *C. difficile* and *B. fragilis* from mono and co-culture during planktonic growth. Data shown is the mean of 3 independent experiments in triplicates and error bars indicate SD, **p < 0.005, ****p<0.0001 as determined by one way ANOVA.

To understand if the inhibitory effect was due to a factor secreted by *B. fragilis*, *C. difficile* and *B. fragilis* were co-cultured under planktonic conditions for 6 h and 10 h (Figure 4D). However, there were no significant differences in *C. difficile* bacterial numbers between mono and co-culture. Additionally, supplementing biofilms with *B. fragilis* planktonic or biofilm culture supernatants did not cause *C. difficile* growth inhibition (Figure S4), indicating that the observed inhibitory effects were specific to adherent biofilms *i.e*. when they are in close proximity to each other.

### LuxS is involved in the *B. fragilis* inhibition of *C. difficile*

To study the role of LuxS in *C. difficile-B. fragilis* interactions, WT *C. difficile* and *luxS* strains were co-cultured with *B. fragilis* in mixed biofilms. CV staining of biofilms showed that there was less *luxS* biofilm formed compared to the WT when cocultured with *B. fragilis* (Figure 5A). While colony counts of *C. difficile* in monocultures were similar for both WT and *luxS* strains (Figure 1B) when cocultured with *B. fragilis* the bacterial counts for both *C. difficile* strains were significantly reduced, although the reduction was significantly higher for the WT than *luxS* (Figure 5B). Colony counts for *B. fragilis* increased significantly during co-culture, with similar levels observed in both co-culture conditions (Figure 5B). These data suggest that AI2/LuxS is involved in mediating the *B. fragilis-induced* inhibition of *C. difficile*, when they are within adherent communities.

**Figure 5.**
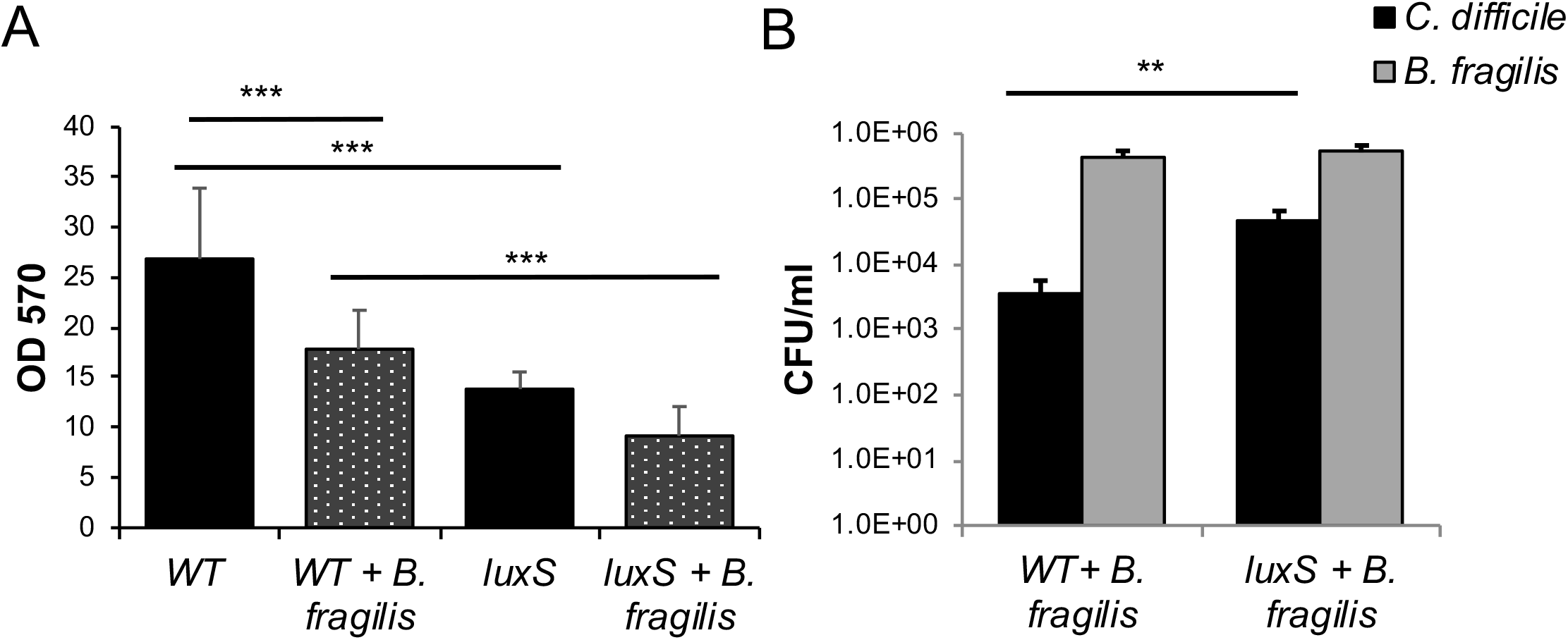
*B. fragilis-mediated* inhibition of *C. difficile* is more prominent for WT than *LuxS*. (**A**) Biofilms for mono and co-cultures of *C. difficile* WT and *luxS* with *B. fragilis* were grown for 24 h and stained with 0.2% CV and were quantified using a spectrophotometer OD_570_. (**B**) Colony counts for *C. difficile* WT, *C. difficile LuxS* during co-culture with *B. fragilis* were performed at 24 h. Data shown is the mean of 3 independent experiments in triplicates and error bars indicate SD, **p < 0.01, ***p<0.001 as determined by one way ANOVA.

### Dual species RNA-seq analysis shows altered metabolism in *C. difficile* and *B. fragilis* in the absence of LuxS

To investigate mechanisms underlying the *C. difficile* inhibition mediated by *B. fragilis*, we performed an RNA-seq analysis to compare biofilm monocultures of *C. difficile* WT, *luxS* or *B. fragilis* with mixed biofilm co-cultures of *C. difficile* WT or *luxS* with *B. fragilis* (Accession number E-MTAB-7523). Differentially expressed genes were defined as having ≥ 1.6-fold relative to their respective control (mono-cultures of either *B. fragilis* or *C. difficile* WT), with an adjusted p-adjusted value ≤ 0.05. A total of 45 genes were up-regulated (21) or down-regulated (24) in *C. difficile* WT (Figure 6, Table 2), while 69 genes were differentially expressed in *C. difficile luxS* of which 34 were down-regulated and 35 up-regulated, during co-culture with *B. fragilis* (Figure 6A, Table 3).

**Figure 6.**
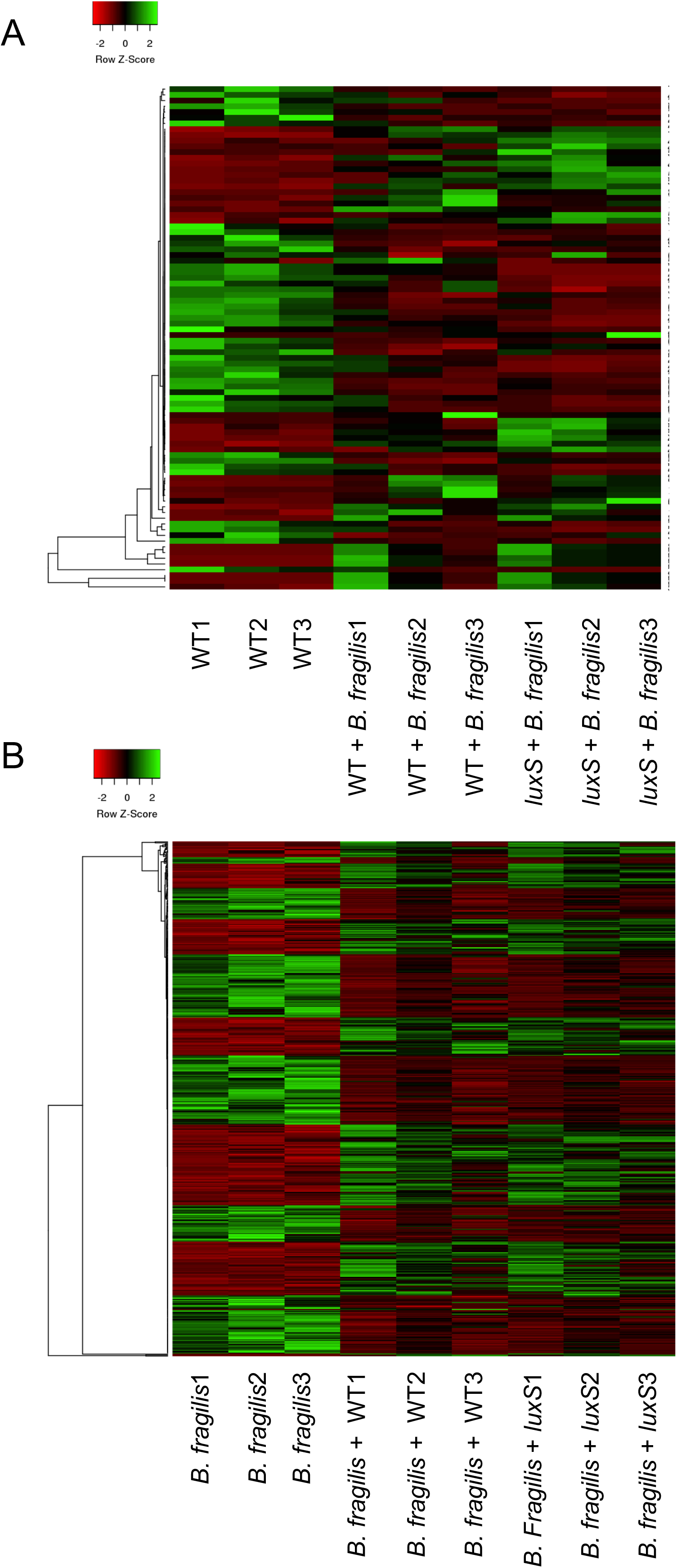
Dual species RNA-seq shows modulation of metabolic pathways in *C. difficile* WT, *luxS* and *B. fragilis*. Heat maps showing clustering of up- and down-regulated genes in (**A**) *C. difficile* WT and *luxS* co-cultured with *B. fragilis* compared to *C. difficile* WT mono-culture, and in (**B**) *B. fragilis* co-cultured with *C. difficile* WT and *luxS* compared to *B. fragilis* monoculture. Red indicates genes that are down-regulated, whilst green indicates genes that are up-regulated.

**Table 2.** Genes up and down-regulated in WT during co-culture with *B. fragilis relative to the* WT *C. difficile* monoculture.

| No | Gene ID | log2FoldChange | Gene Annotation |
| --- | --- | --- | --- |
| 1 | CDR20291_0194 | 1.3822081 | 10 kDa chaperonin |
| 2 | CDR20291_1016 | 1.7087847 | glycerol-3-phosphate acyltransferase PlsX |
| 3 | CDR20291_1017 | 1.7011992 | 3-oxoacyl-ACP synthase III |
| 4 | CDR20291_1018 | 1.2469326 | trans-2-enoyl-ACP reductase |
| 5 | CDR20291_1337 | 1.3290499 | transcriptional regulator |
| 6 | CDR20291_1861 | 1.412304 | biotin carboxylase acetyl-CoA carboxylase subunit A |
| 7 | CDR20291_2027 | 1.6309458 | 2-nitropropane dioxygenase |
| 8 | CDR20291_3225 | 1.0229025 | formate/nitrite transporter |
| 9 | CDR20291_0363 | -1.377088 | radical SAM protein |
| 10 | CDR20291_0364 | -1.003595 | hypothetical protein |
| 11 | CDR20291_0365 | -1.461207 | (R)-2-hydroxyisocaproate dehydrogenase |
| 12 | CDR20291_0659 | -1.248536 | radical SAM protein |
| 13 | CDR20291_1271 | -1.15672 | hypothetical protein |
| 14 | CDR20291_1309 | -1.208811 | phosphohydrolase |
| 15 | CDR20291_1400 | -1.833327 | imidazole glycerol phosphate synthase subunit HisH |
| 16 | CDR20291_1834 | -1.411634 | ethanolamine/propanediol ammonia-lyase heavy chain |
| 17 | CDR20291_2416 | -1.231651 | hypothetical protein |
| 18 | CDR20291_2417 | -1.574982 | hypothetical protein |
| 19 | CDR20291_2610 | -1.046807 | two-component sensor histidine kinase |

Note: P-adjusted value $\leq 0.05$

**Table 3.** Genes up and down-regulated in *luxS* during co-culture with *B. fragilis relative to the* WT *C. difficile* monoculture.

| No | Gene ID | log2FoldChange | Gene Annotation |
| --- | --- | --- | --- |
| 1 | CDR20291_0491 | 1.0964355 | RNA methylase |
| 2 | CDR20291_0492 | 1.1802034 | hypothetical protein |
| 3 | CDR20291_0493 | 1.0076105 | outer membrane lipoprotein |
| 4 | CDR20291_0715 | 1.6245725 | N-acetylmuramoyl-L-alanine amidase |
| 5 | CDR20291_1366 | 0.9274027 | ferrous ion transport protein |
| 6 | CDR20291_1374 | 1.1651176 | iron-sulfur protein |
| 7 | CDR20291_1691 | 1.4187304 | nitrite and sulfite reductase subunit |
| 8 | CDR20291_1716 | 1.3143654 | thiol peroxidase |
| 9 | CDR20291_1717 | 1.110872 | hypothetical protein |
| 10 | CDR20291_1934 | 1.5456229 | hypothetical protein |
| 11 | CDR20291_1936 | 1.3368874 | GntR family transcriptional regulator |
| 12 | CDR20291_1937 | 1.3736945 | ABC transporter ATP-binding protein |
| 13 | CDR20291_2389 | 1.2619709 | competence protein |
| 14 | CDR20291_2830 | 1.11333 | ribonucleoside-diphosphate reductase subunit alpha |
| 15 | CDR20291_2928 | 1.7538156 | PTS system transporter subunit IIABC |
| 16 | CDR20291_2930 | 1.5713168 | trehalose-6-phosphate hydrolase |
| 17 | CDR20291_3075 | 1.3649687 | osmoprotectant ABC transporter substrate-binding/permease |
| 18 | CDR20291_3104 | 0.8099098 | sigma-54-dependent transcriptional activator |
| 19 | CDR20291_3434 | 1.4402419 | homocysteine S-methyltransferase |
| 20 | CDR20291_0025 | -1.32053 | acetoin:2%2C6-dichlorophenolindophenol oxidoreductase subunit alpha |
| 21 | CDR20291_0615 | -0.86915 | nucleotide phosphodiesterase |
| 22 | CDR20291_0802 | -2.260047 | ABC transporter substrate-binding protein |
| 23 | CDR20291_0911 | -1.24181 | electron transfer flavoprotein subunit beta |
| 24 | CDR20291_1359 | -0.850952 | hypothetical protein |
| 25 | CDR20291_1370 | -1.124726 | tyrosyl-tRNA synthetase |
| 26 | CDR20291_1497 | -1.456296 | phosphomethylpyrimidine kinase |
| 27 | CDR20291_1498 | -1.727285 | hydroxyethylthiazole kinase |
| 28 | CDR20291_1499 | -1.276387 | thiamine-phosphate pyrophosphorylase |
| 29 | CDR20291_1591 | -1.51799 | dinitrogenase iron-molybdenum cofactor |
| 30 | CDR20291_1901 | -1.832663 | ABC transporter ATP-binding protein |
| 31 | CDR20291_1902 | -1.875507 | ABC transporter substrate-binding protein |
| 32 | CDR20291_1903 | -1.659329 | ABC transporter permease |
| 33 | CDR20291_1904 | -1.638609 | hypothetical protein |
| 34 | CDR20291_1925 | -1.54806 | flavodoxin |
| 35 | CDR20291_2474 | -1.434272 | DNA-directed RNA polymerase subunit omega |
| 36 | CDR20291_2515 | -1.603951 | amino acid permease family protein |
| 37 | CDR20291_2516 | -1.176482 | cobalt dependent x-pro dipeptidase |
| 38 | CDR20291_2660 | -0.958814 | teichuronic acid biosynthesis glycosyl transferase |
| 39 | CDR20291_2870 | -2.335518 | hypothetical protein |
| 40 | CDR20291_3142 | -1.665916 | pyrroline-5-carboxylate reductase |
| 41 | CDR20291_3143 | -1.830481 | formate acetyltransferase |
| 42 | CDR20291_3144 | -1.643573 | pyruvate formate-lyase 3 activating enzyme |

Note: P-adjusted value $\leq 0.05$

Eight up-regulated genes were specific to *C. difficile* WT co-culture (Table 2). Of these, four genes involved in fatty acid biosynthesis and metabolism: *fabH* encoding 3-oxoacyl-[acyl-carrier protein] synthase III, *fabK* encoding trans-2-enoyl-ACP reductase, *accC* encoding a biotin carboxylase (acetyl-CoA carboxylase subunit A), and *accB* encoding a biotin carboxyl carrier protein of acetyl-CoA carboxylase (Table 2). 11 genes were down-regulated exclusively in WT however, these genes do not coincide with a specific metabolic pathway.

18 up-regulated genes were specific to *luxS* in co-culture (Table 3). These include a putative homocysteine S-methyltransferase, a putative osmoprotectant ABC transporter, substrate binding/ permease protein, ribonucleoside-diphosphate reductase alpha chain (*nrdE*) and two genes from the trehalose operon: a PTS system II ABC transporter, and trehalose-6-phosphate hydrolase (treA). 24 genes were down-regulated (Table 3), which include 3 genes involved in thiamine metabolism *thiD*, *thiK* and *thiE1*, (CDR20291_1497, CDR20291_1498 and CDR20291_1499 respectively) which encode a putative phosphomethylpyrimidine kinase, 4-methyl-5-beta-hydroxyethylthiazole kinase and thiamine-phosphate pyrophosphorylase respectively.

A total of 26 genes were differentially expressed in both *C. difficile* WT and *C. difficile luxS* when co-cultured with *B. fragilis* (Figure 6A, Table 4). These include six up-regulated genes (*accB*, *abfH*, *abfT*, *abfD*, *sucD* and *cat1*) involved in carbon and butanoate metabolism, with *cat1*, which encodes succinyl-CoA:coenzyme A transferase, being the highest up-regulated gene for both *C. difficile* strains.

**Table 4.** *Genes up- and down-regulated in* both WT and *luxS* during co-culture with *B. fragilis relative to the* WT *C. difficile* monoculture.

| No | Gene ID | Log2 Foldchange | Gene Annotation |
| --- | --- | --- | --- |
| 1 | CDR20291_0511 | -1.947939847 | hypothetical protein |
| 2 | CDR20291_0512 | -1.790659218 | hypothetical protein |
| 3 | CDR20291_0910 | -1.118039675 | butyryl-CoA dehydrogenase |
| 4 | CDR20291_1799 | -1.559955287 | hypothetical protein |
| 5 | CDR20291_1878 | -1.084562908 | ABC transporter ATP-binding protein |
| 6 | CDR20291_2418 | -1.624152909 | hypothetical protein |
| 7 | CDR20291_2419 | -1.384788131 | aminotransferase |
| 8 | CDR20291_2611 | -1.45763049 | two-component response regulator |
| 9 | CDR20291_2612 | -1.940640211 | hypothetical protein |
| 10 | CDR20291_2613 | -1.569696353 | polysaccharide deacetylase |
| 11 | CDR20291_0196 | 1.242760946 | hypothetical protein |
| 12 | CDR20291_0197 | 1.024719474 | hypothetical protein |
| 13 | CDR20291_1015 | 1.855396639 | fatty acid biosynthesis transcriptional regulator |
| 14 | CDR20291_1862 | 1.571375262 | biotin carboxyl carrier protein of acetyl-CoA carboxylase |
| 15 | CDR20291_1931 | 1.501260024 | AraC family transcriptional regulator |
| 16 | CDR20291_1935 | 1.754871647 | hypothetical protein |
| 17 | CDR20291_2077 | 1.310107407 | Sodium:dicarboxylate symporter |
| 18 | CDR20291_2140 | 2.555234046 | N-acetylmannosamine-6-phosphate 2-epimerase |
| 19 | CDR20291_2227 | 3.630122723 | NAD-dependent 4-hydroxybutyrate dehydrogenase |
| 20 | CDR20291_2228 | 3.789825947 | 4-hydroxybutyrate CoA transferase |
| 21 | CDR20291_2229 | 3.695347944 | hypothetical protein |
| 22 | CDR20291_2230 | 3.544168193 | gamma-aminobutyrate metabolism dehydratase/isomerase |
| 23 | CDR20291_2231 | 3.864728796 | succinate-semialdehyde dehydrogenase [NAD(P)+] |
| 24 | CDR20291_2232 | 4.2239099 | succinyl-CoA:coenzyme A transferase |
| 25 | CDR20291_2233 | 4.339535053 | hypothetical protein |
| 26 | CDR20291_3110 | 1.176093028 | hypothetical protein |

Note: P-adjusted value $\leq 0.05$

In contrast, a higher number of genes (266) were differentially expressed in *B. fragilis* when co-cultured with *C. difficile* (WT and *luxS*) (Table S3, Figure 6B). A total of 114 *B. fragilis* genes were found to be specific to *C. difficile* WT co-culture, with 56 of these up-regulated and 58 down-regulated (Table 5). Similarly, 91 genes were found to be specific to *C. difficile luxS* co-culture, with 56 of these up-regulated and 35 down-regulated (Table 6). Although distinct *B. fragilis* expression profiles were observed with the WT and *luxS* (Figure 6B), there were no clear pathways identified in the datasets.

**Table 5.**
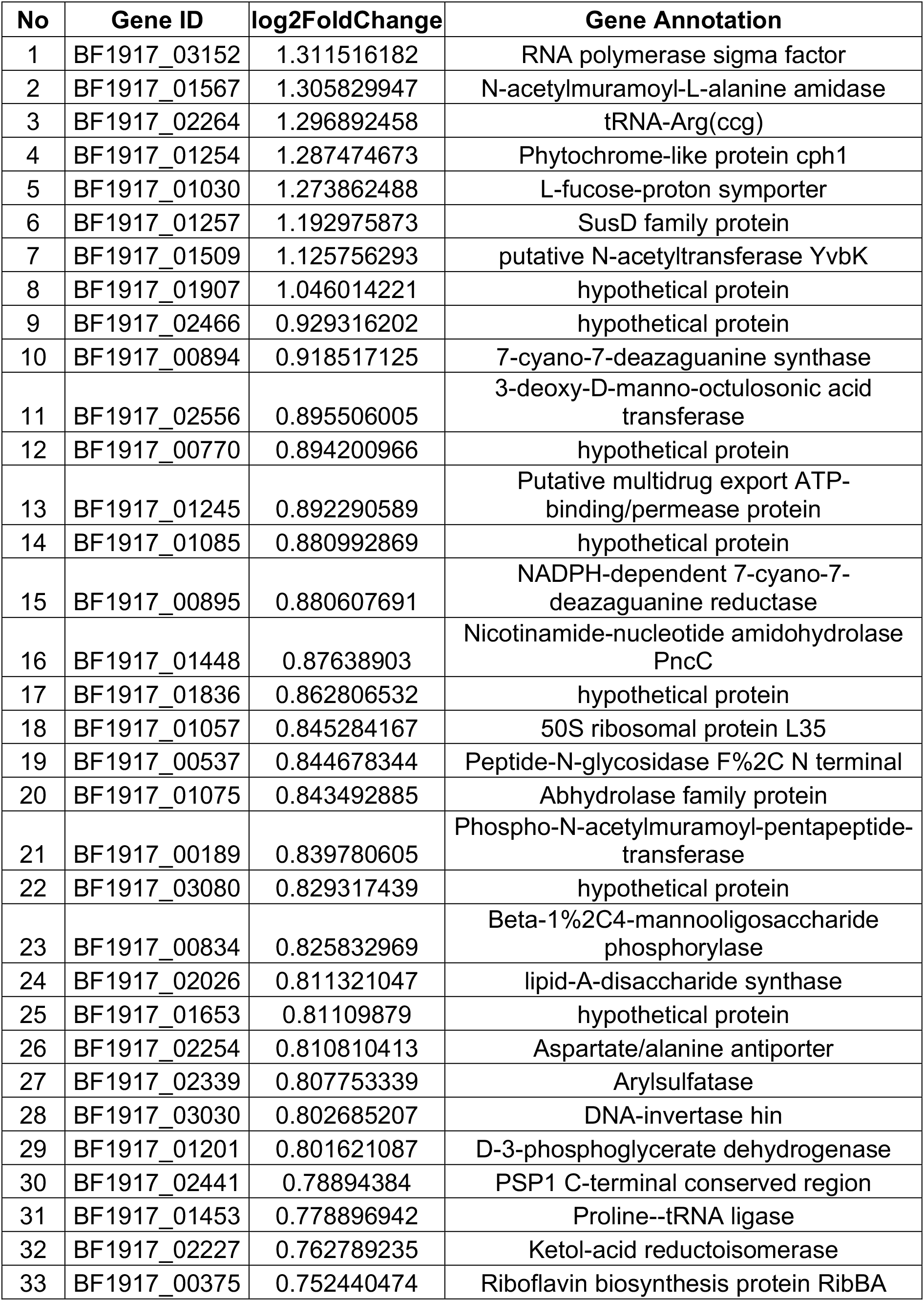
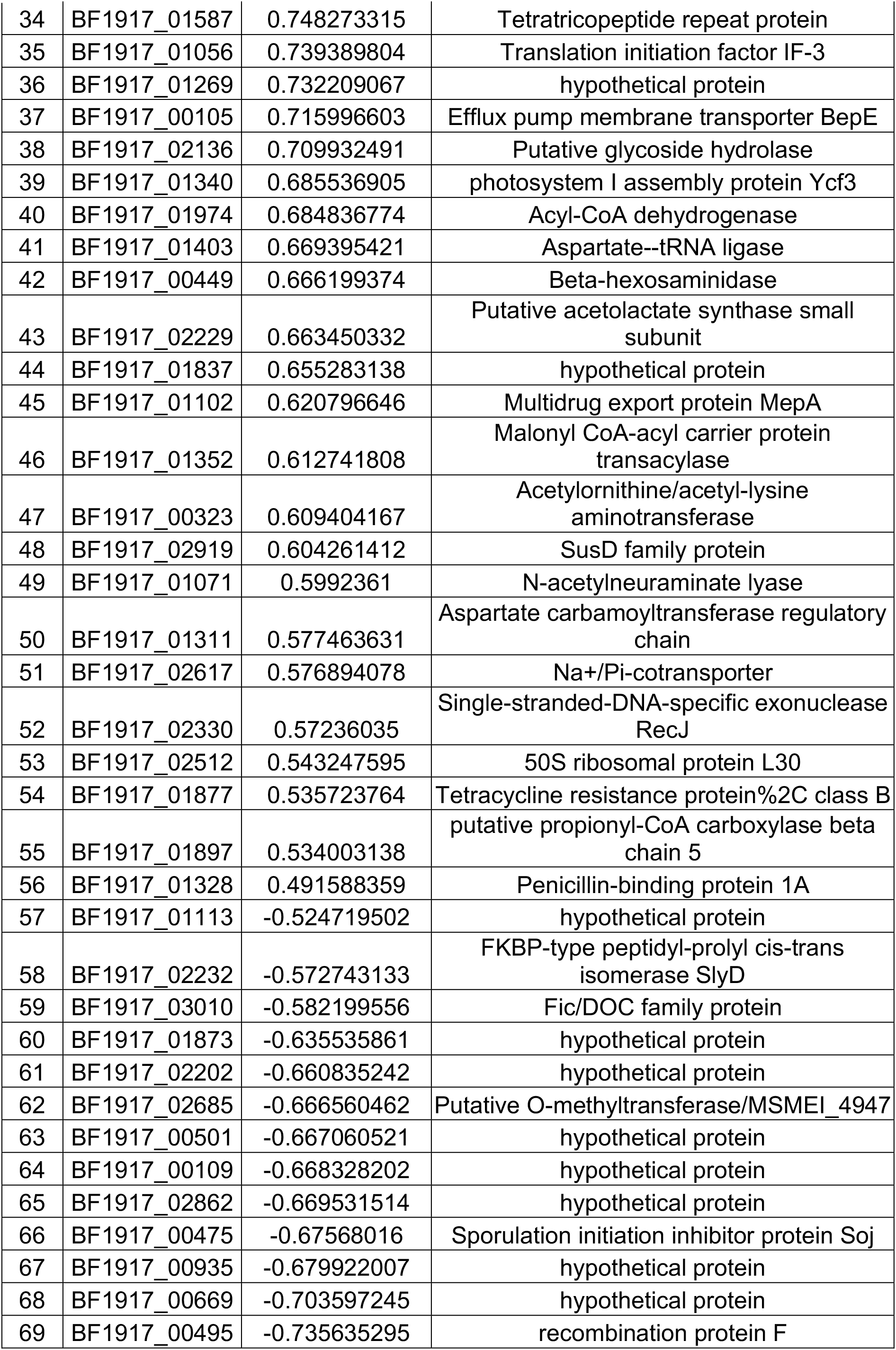
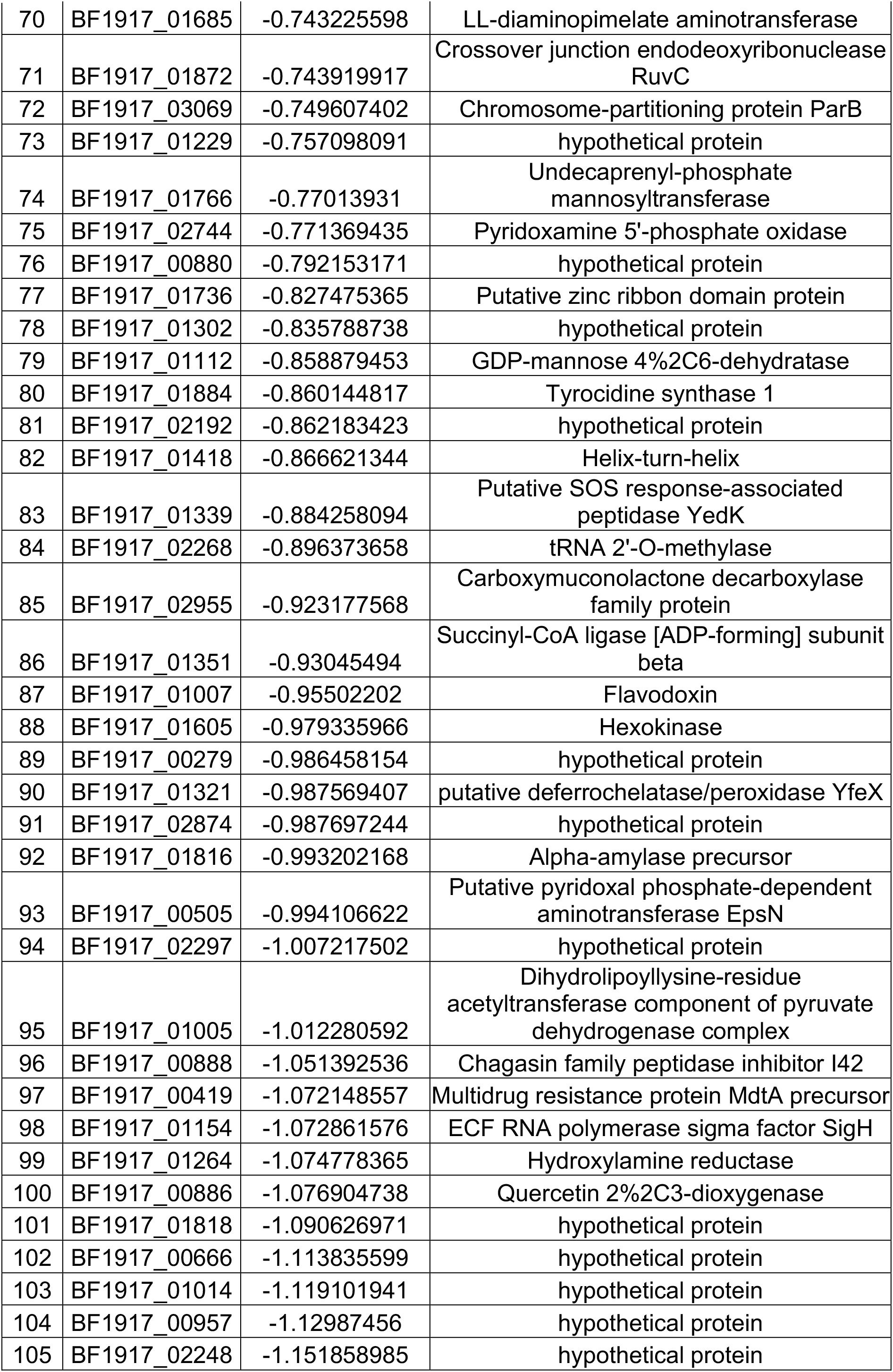
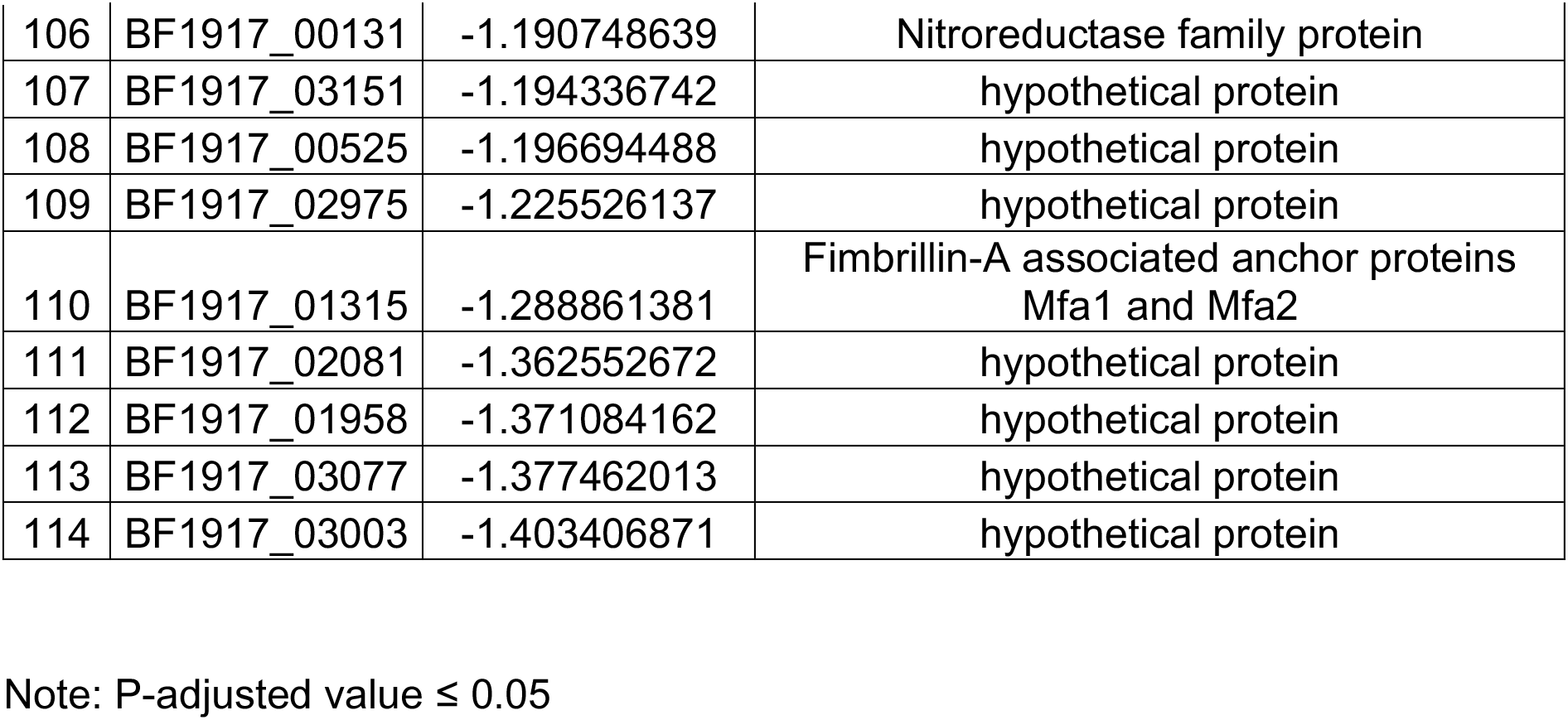
Genes up- and down-regulated *in B. fragilis* specific to co-culture with WT *C. difficile*.

**Table 6.** Genes up- and down-regulated *in B. fragilis* specific to co-culture with *luxS C. difficile*.

| No | Gene ID | log2FoldChange | Gene Annotation |
| --- | --- | --- | --- |
| 1 | BF1917_03116 | 1.50249 | hypothetical protein |
| 2 | BF1917_00412 | 1.48934 | Tyrosine recombinase XerD |
| 3 | BF1917_02127 | 1.295481 | fec operon regulator FecR |
| 4 | BF1917_02593 | 1.280656 | tRNA-Leu(caa) |
| 5 | BF1917_01792 | 1.235375 | Reverse rubrerythrin-1 |
| 6 | BF1917_02909 | 1.220847 | hypothetical protein |
| 7 | BF1917_01427 | 1.200963 | ATP-dependent Clp protease ATP-binding subunit ClpC |
| 8 | BF1917_02997 | 1.188287 | hypothetical protein |
| 9 | BF1917_01065 | 1.176729 | tRNA-Thr(cgt) |
| 10 | BF1917_00574 | 1.133713 | hypothetical protein |
| 11 | BF1917_01993 | 1.109684 | 60 kDa chaperonin |
| 12 | BF1917_01994 | 1.09403 | 10 kDa chaperonin |
| 13 | BF1917_02911 | 1.08306 | Tyrosine recombinase XerC |
| 14 | BF1917_02143 | 1.066955 | NAD-specific glutamate dehydrogenase |
| 15 | BF1917_02972 | 1.063195 | fec operon regulator FecR |
| 16 | BF1917_00620 | 1.03149 | fec operon regulator FecR |
| 17 | BF1917_01491 | 1.021451 | hypothetical protein |
| 18 | BF1917_02039 | 1.015769 | tRNA-Phe(gaa) |
| 19 | BF1917_03113 | 0.990297 | hypothetical protein |
| 20 | BF1917_03172 | 0.98787 | tRNA-Arg(gcg) |
| 21 | BF1917_00358 | 0.980119 | MarR family protein |
| 22 | BF1917_00832 | 0.956639 | hypothetical protein |
| 23 | BF1917_00767 | 0.951389 | hypothetical protein |
| 24 | BF1917_01889 | 0.938171 | Septum formation protein Maf |
| 25 | BF1917_01282 | 0.932255 | ATP synthase subunit beta |
| 26 | BF1917_03056 | 0.92593 | hypothetical protein |
| 27 | BF1917_02999 | 0.921887 | hypothetical protein |
| 28 | BF1917_00178 | 0.905375 | 30S ribosomal protein S20 |
| 29 | BF1917_02814 | 0.89716 | hypothetical protein |
| 30 | BF1917_02866 | 0.885735 | hypothetical protein |
| 31 | BF1917_02896 | 0.871156 | hypothetical protein |
| 32 | BF1917_00295 | 0.865619 | 50S ribosomal protein L34 |
| 33 | BF1917_03169 | 0.843942 | tRNA-Asp(gtc) |
| 34 | BF1917_01072 | 0.843727 | Cellobiose 2-epimerase |
| 35 | BF1917_02507 | 0.835563 | 50S ribosomal protein L36 |
| 36 | BF1917_02553 | 0.817012 | 30S ribosomal protein S21 |
| 37 | BF1917_03177 | 0.802377 | tRNA-Lys(ttt) |
| 38 | BF1917_01910 | 0.800503 | 50S ribosomal protein L31 type B |
| 39 | BF1917_01181 | 0.795826 | putative ABC transporter ATP-binding protein YjjK |
| 40 | BF1917_03168 | 0.776641 | tRNA-Pro(tgg) |
| 41 | BF1917_02950 | 0.740702 | transcriptional regulator BetI |
| 42 | BF1917_02033 | 0.735701 | hypothetical protein |
| 43 | BF1917_01372 | 0.732434 | hypothetical protein |
| 44 | BF1917_00179 | 0.697125 | tRNA-Glu(ttc) |
| 45 | BF1917_01819 | 0.691297 | 4Fe-4S binding domain protein |
| 46 | BF1917_01081 | 0.670376 | F5/8 type C domain protein |
| 47 | BF1917_01847 | 0.651964 | hypothetical protein |
| 48 | BF1917_02253 | 0.635549 | Methylmalonyl-CoA mutase |
| 49 | BF1917_01261 | 0.623302 | hypothetical protein |
| 50 | BF1917_02778 | 0.561439 | enterobactin exporter EntS |
| 51 | BF1917_02391 | 0.556052 | 50S ribosomal protein L13 |
| 52 | BF1917_02815 | 0.541997 | Phosphoenolpyruvate carboxykinase [ATP] |
| 53 | BF1917_02820 | 0.538608 | Phosphoenolpyruvate carboxykinase [ATP] |
| 54 | BF1917_02819 | 0.535546 | Phosphoenolpyruvate carboxykinase [ATP] |
| 55 | BF1917_02049 | 0.508167 | putative enoyl-CoA hydratase 1 |
| 56 | BF1917_00123 | 0.446281 | Acyl carrier protein |
| 57 | BF1917_02261 | -0.51363 | Glucose-6-phosphate isomerase |
| 58 | BF1917_00321 | -0.53822 | DNA-binding transcriptional repressor PuuR |
| 59 | BF1917_00914 | -0.56999 | Bifunctional transcriptional activator/DNA repair enzyme AdaA |
| 60 | BF1917_00594 | -0.60044 | DNA replication and repair protein RecF |
| 61 | BF1917_01306 | -0.60418 | Formate--tetrahydrofolate ligase |
| 62 | BF1917_00287 | -0.64751 | Phosphate acetyltransferase |
| 63 | BF1917_00206 | -0.68829 | tetratricopeptide repeat protein |
| 64 | BF1917_00107 | -0.72261 | Beta-galactosidase |
| 65 | BF1917_01926 | -0.72606 | Outer membrane efflux protein |
| 66 | BF1917_01224 | -0.7469 | putative GTP-binding protein EngB |
| 67 | BF1917_01293 | -0.75882 | Transcriptional regulatory protein CusR |
| 68 | BF1917_01212 | -0.76206 | hypothetical protein |
| 69 | BF1917_01767 | -0.77069 | Dihydroorotase |
| 70 | BF1917_01707 | -0.78791 | hypothetical protein |
| 71 | BF1917_00291 | -0.79844 | UDP-2%2C3-diacylglucosamine hydrolase |
| 72 | BF1917_01301 | -0.80259 | dTDP-4-dehydrorhamnose reductase |
| 73 | BF1917_02606 | -0.8053 | Aminopeptidase YwaD precursor |
| 74 | BF1917_01852 | -0.81525 | Transaldolase |
| 75 | BF1917_01735 | -0.83554 | putative oxidoreductase |
| 76 | BF1917_02275 | -0.8445 | FMN reductase [NAD(P)H] |
| 77 | BF1917_02683 | -0.86216 | Lipoprotein-releasing system<br>transmembrane protein LolE |
| 78 | BF1917_00953 | -0.87106 | putative A/G-specific adenine glycosylase<br>YfhQ |
| 79 | BF1917_00810 | -0.9142 | hypothetical protein |
| 80 | BF1917_02167 | -0.91437 | Pyruvate formate-lyase 1-activating enzyme |
| 81 | BF1917_02588 | -0.9256 | Bifunctional transcriptional activator/DNA<br>repair enzyme Ada |
| 82 | BF1917_00809 | -0.98589 | Cadmium%2C zinc and cobalt-transporting<br>ATPase |
| 83 | BF1917_01476 | -1.01713 | Colicin I receptor precursor |
| 84 | BF1917_01210 | -1.01929 | putative metallo-hydrolase |
| 85 | BF1917_00766 | -1.03246 | hypothetical protein |
| 86 | BF1917_01961 | -1.06955 | Molecular chaperone Hsp31 and glyoxalase<br>3 |
| 87 | BF1917_02112 | -1.11145 | Lactate utilization protein C |
| 88 | BF1917_00727 | -1.19678 | hypothetical protein |
| 89 | BF1917_03146 | -1.33023 | hypothetical protein |
| 90 | BF1917_01478 | -1.33473 | hypothetical protein |
| 91 | BF1917_02716 | -1.4039 | hypothetical protein |

Note: P-adjusted value $\leq 0.05$

Whilst the highest up-regulated gene in both WT and LuxS co-cultures encodes a putative virus attachment protein, no other viral genes were shown to be up-regulated (Table S3). Genes encoding iron containing proteins desulfoferrodoxin and rubrerythrin were highly up-regulated in both conditions. Interestingly, multiple copies of *fecR*, a key regulator for the ferric citrate transport system [50], and ferrous iron transport protein B were also up-regulated. Additionally, a number of other metabolic pathways were up-regulated, including four genes (encoding 3-isopropylmalate dehydratase small subunit, 3-isopropylmalate dehydrogenase, Galactokinase and 3-isopropylmalate dehydratase large subunit) involved in valine, leucine and isoleucine biosynthesis and C5-branched dibasic acid metabolism. It should be noted that many of the up-regulated genes were hypothetical proteins of unknown function. Similarly, several metabolic pathways were down-regulated in both co-culture conditions.

These include six genes involved in carbon metabolism, four genes involved in alanine, aspartate and glutamate metabolism, and four genes involved in the biosynthesis of amino acids, although these appear to be single genes rather than specific pathways.

## Discussion

Inter-bacterial interactions within gut communities are critical in controlling invasion by intestinal pathogens. Quorum sensing molecules such as AI-2 are instrumental in bacterial communication, especially during formation of bacterial communities [32, 37, 38, 51-54]. *C. difficile* produces AI-2, although the mechanism of action of LuxS/AI-2 in *C. difficile*, particularly within a biofilm community is unclear. We report here that the *C. difficile* LuxS/AI-2 plays an important role in the formation of single and multi-species communities. In *C. difficile* biofilms, LuxS mediates the induction of prophages, which likely contributes to the biofilm structure. Whereas, in a mixed biofilm of *C. difficile* and the intestinal commensal and pathogen, *B. fragilis*, LuxS likely triggers the induction of differential metabolic responses in *B. fragilis*, that leads to growth inhibition of *C. difficile*. To our knowledge, this is the first time dual species RNA-seq [55] has been applied to analyse interactions between anaerobic gut bacteria in an adherent biofilm community.

Bacterial biofilms contain a number of extracellular components that make up their complex structure including extracellular DNA (eDNA), a key component that binds together bacteria within a community. Autolysis is a common mechanism by which eDNA is released from bacterial cells [56]. In bacteria such as *Staphylococcus aureus* and *Pseudomonas aeruginosa*, eDNA is generated through the lysis of subpopulations within a biofilm, under the control of quorum sensing [56-59]. In *C. difficile luxS* mutant (*luxS*) biofilms, we observe reduced induction of two *C. difficile* prophages compared with the WT. These phage loci were conserved in several *C. difficile* strains with the Region 2 encoding a phiC2-like phage [60]. Given that it has previously been shown that eDNA is a major component of *C. difficile* biofilms [27, 61], it is likely that phage-mediated bacterial cell lysis and subsequent DNA release help build a biofilm. However, phages are also known to control biofilm structure in some organisms: a filamentous phage of *P. aeruginosa* was reported to be a structural component of the biofilm [62] and an AI-2 induced phage mediated the dispersal of *Enterococcus faecalis* biofilms [43]. Although attempts to visualise phages from *C. difficile* biofilms with transmission electron microscopy (data not shown) were unsuccessful, we cannot rule out the possibility that *C. difficile* phages may directly influence the biofilm structure. While the precise mechanisms by which LuxS/AI-2 controls phage induction are yet to be elucidated, AI-2 appears to be signalling through a yet unidentified AI-2 receptor in *C. difficile*. Phage-mediated control of biofilms may in part explain the variation observed in biofilm formation between different *C. difficile* strains [63].

The human gut hosts a variety of bacterial species, which compete or coexist with each other. It is likely that the bacteria occupying this niche form multi-species bacterial communities in association with the mucus layer. Interactions within such communities are important in gaining a better understanding of phenomena such as ‘colonisation resistance’ which prevents pathogens such as *C. difficile* from establishing an infection [64]. Whilst sequencing studies have identified members of the *Bacteroides* genus as being associated with gut colonisation resistance to *C. difficile*, the mechanisms have remained elusive [10]. A recent study demonstrated that production of the enzyme: bile salt hydrolase, is responsible for the inhibitory effect of *B. ovatus* on *C. difficile* [65]. This study reported that in the presence of bile acids, cell free supernatants for *B. ovatus* were capable of inhibiting the growth of *C. difficile* whereas in the absence of bile acids, *C. difficile* growth was promoted [65]. Since bile acids are not supplemented into our media, a different mechanism is likely responsible for *B. fragilis* mediated inhibition of *C. difficile*. Also, the growth restraining effects of *B. fragilis* on *C. difficile* were evident only within mixed biofilms, not in planktonic culture or with culture supernatants. While it is likely that cell-cell contact is essential for the inhibitory effect, we cannot exclude involvement of an inhibitory secreted molecule that accumulates to a higher concentration within a biofilm environment, or that *B. fragilis* has a competitive growth advantage in a biofilm environment.

A dual species RNA-seq analysis performed to understand the interactions between the two bacterial species, showed that largely all the differentially expressed genes mapped to distinct metabolic pathways. Overall, a higher number of genes were modulated in *B. fragilis* as compared to *C. difficile* strain during co-culture, which is in line with the growth characteristics observed. Carbon and butanoate metabolism pathways were induced in *C. difficile* strains in response to co-culture (*accB*, *abfH*, *abfT*, *abfD*, *sucD* and *cat1*). As *B. fragilis* is known to produce succinate [66], it is likely that the upregulation in these pathways results from the increased levels of succinate in the culture medium. However, since gut microbiota-produced succinate promotes *C. difficile* growth *in vivo* [17], it is unlikely that these changes are directly responsible for the observed inhibition of *C. difficile*. However, bacteria utilise carbohydrates in a sequential manner [67]. Consistent with this, we observed a down-regulation of genes important for the utilisation of pyruvate such as *bcd2* and *idhA* encoding for butyryl-CoA dehydrogenase and (r)-2-hydroxyisocaproate dehydrogenase respectively. Such a shift in metabolism could allow *B. fragilis* to fully consume other metabolites, and thus enabling it to outcompete *C. difficile*.

Additionally, it is interesting to note that a number of copies of the ferric citrate transport system regulator, *fecR*, are up-regulated in *B. fragilis* during *C. difficile* coculture. The ferric citrate transport system is an iron uptake system that responds to the presence of citrate [50, 68, 69]. Analysis of the *C. difficile* genome using BLAST (NCBI) showed that *C. difficile* does not possess this iron uptake system. Given the evidence that ferric citrate is an iron source in the gut [50, 70], *B. fragilis* may have an advantage over *C. difficile* in sequestering iron, and thus preventing *C. difficile* colonisation. Although the clear modulation of metabolic pathways strongly suggest a competitive advantage of *B. fragilis* over *C. difficile*, it is possible that the genes with unknown functions that are differentially expressed in *B. fragilis* (Table S1), encode pathways for the production of yet to be identified small inhibitory molecules. The LuxS/AI-2 quorum sensing system is known to have a cross-species signalling role in many bacteria [48, 71, 72]. While all sequenced *C. difficile* strains produce AI-2, only selected strains of *B. fragilis* have the ability to produce AI-2 [73]. A recent study showed that *Ruminococcus obeuem* inhibited *Vibrio cholerae* in the gut via LuxS/AI-2 mediated downregulation of *V. cholerae* colonisation factors [74]. Also, AI-2 produced by engineered *E. coli* was reported to influence firmicutes/bacteroidetes ratios in microbiota treated with streptomycin [75]. Our data show the involvement of LuxS/AI-2 in the *B. fragilis-mediated C. difficile* growth inhibition. As the *B. fragilis* strain used in this study does not produce AI-2, it is likely that *B. fragilis* responds differentially to AI-2 produced by *C. difficile*. Similar to the WT, most transcriptional changes were in metabolic pathways, although specific sets of genes were modulated in the *luxS* mutant. Modulation of prophage genes was not seen, unlike single biofilm cultures, indicating different dominant mechanisms at play in a multispecies environment.

It was interesting to note that the trehalose utilisation operon, which provides a growth advantage to *C. difficile* against other gut bacteria [76], was upregulated in both the single *luxS* biofilms (Table 1) and *luxS* co-cultured with *B. fragilis* (Table 3). The upregulation of a phosphotransferase system component and *treA* (both in the same operon), likely enables increased utilisation of trehalose, providing an additional carbon source. Like glucose, trehalose acts as an osmoprotectant [77] and its presence within the cell may help maintain protein conformation during cellular dehydration. It is possible that trehalose plays a role in building *C. difficile* biofilms, as reported for *Candida* biofilms [78]. Our preliminary studies with exogenous trehalose levels similar to those used by Collins *et al*. (2018) showed an inhibitory effect on biofilm formation by both WT and *luxS*, although no differential effects were observed between *luxS* and WT (data not shown). However, further investigations into accumulation of trehalose within biofilms are required to clarify the role of trehalose in *luxS* mediated biofilm formation.

In conclusion, we report that *C. difficile* LuxS/AI-2 may play a key role in building *C. difficile* communities through mediating prophage induction, and subsequent accumulation of eDNA. In mixed communities, *C. difficile* AI-2 likely signals to *B. fragilis* to induce an altered metabolic response, enabling it to outgrow *C. difficile*. Further studies are required to understand the precise AI-2 sensing pathways involved.

## Materials and Methods

### Bacterial strains and media

Two bacterial species were used in this study – *C. difficile* strain: B1/NAP1/027 R20291 (isolated from the Stoke Mandeville outbreak in 2004 and 2005), and *Bacteroides fragilis* (kindly provided by Dr Gianfranco Donelli, Rome). A luxS Clostron R20291 mutant described previously in Dapa et al, 2013 was used in this study. Both species were cultured under anaerobic conditions (80% N_2_, 10% CO_2_, 10% H_2_) at 37°C in an anaerobic workstation (Don Whitley, United Kingdom) in BHIS, supplemented with L-Cysteine (0.1% w/v; Sigma, United Kingdom), yeast extract (5 g/l; Oxoid) and glucose (0.1 M).

*Vibrio harveyi* strain: BB170 was used to measure AI-2. *V. harveyi* strains were cultured in aerobic conditions at 30°C in Lysogeny broth (LB) supplemented with kanamycin (50 μg/ml).

### Biofilm formation assay

Biofilms were grown as per the previously published protocol [27]. Overnight cultures of *C. difficile* were diluted 1:100 in fresh BHIS with 0.1 M glucose. 1 ml aliquots were pipetted into 24-well tissue culture treated polystyrene plates (Costar), and incubated under anaerobic condition at 37°C, for 6 - 120 h. Tissue culture plates were pre-incubated for 48h prior to use. The plates were wrapped with parafilm to prevent liquid evaporation.

### Measurement of biofilm biomass

Biofilm biomass was measured using crystal violet (CV) [27, 79]. After the required incubation, each well of the 24-well plate was washed with sterile phosphate buffer saline (PBS) and allowed to dry for a minimum of 10 mins. The biofilm was stained using 1 ml of filter-sterilised 0.2% CV and incubated for 30 mins at 37°C, in anaerobic conditions. The CV was removed from each well, and wells were subsequently washed twice with sterile PBS. The dye was extracted by incubated with 1 ml methanol for 30 mins at room temperature (RT) in aerobic conditions. The methanol-extracted dye was diluted 1:1, 1:10 or 1:100 and OD_570_ was measured with a spectrophotometer (Biochrom, UK).

For bacterial cell counts from the biofilm, the planktonic phase was removed and wells were washed once using sterile PBS. The adherent biofilms were then detached by scrapping with a sterile pipette tip and re-suspended into 1 ml PBS. Serial dilutions were made and plated onto BHIS plates to determine the CFU present in the biofilm.

### Co-culture biofilm assay

For generation of co-culture biofilms, both *C. difficile* and B. *fragilis* were diluted to an OD_600_ of 1. Both species were diluted 1:100 into fresh BHIS with 0.1 M glucose. Biofilms assays were performed as described above and measured by a combination of CV staining and CFU. To distinguish between *C. difficile* and *B. fragilis*, serial dilutions used for determining CFU were plated on BHIS plates additionally supplemented with *C. difficile* selective supplement (Oxoid, UK). Colonies can be differentiated by size and colony morphology as *B. fragilis* form very small colonies.

### Exogenous addition of DPD

To analyse the potential signalling role of AI-2, biofilm assays were performed as described above in BHIS with 0.1 M glucose containing 1 nM, 10 nM, 100 nM, or 1 μM of chemically synthesised, exogenous 4,5-Dihydroxy-2,3-pentanedione (Omm Scientific, Texas USA) for both *C. difficile* WT and LuxS. BHIS with 0.1 M glucose was used as a control. Samples were washed and stained with 0.2% CV at either 24 h or 72 h.

### AI-2 Assay

The AI-2 bioluminescence assay was carried out essentially as described by Bassler *et al*. 1993 [80]. The *V. harveyi* reporter strain BB170 was grown overnight in LB medium before being diluted 1: 5000 in Autoinducer Bioassay (AB) medium containing 10 % (v/v) cell-free conditioned medium collected from either planktonic or biofilm *C. difficile* cultures (in BHI) and allowed to grow at 30°C with shaking. AB medium containing 10 % (v/v) from *V. harveyi* BB120 was used as a positive control, and 10 % (v/v) sterile BHI medium as a blank. Luminescence was measured every hour using a SPECTROstar Omega plate reader. Induction of luminescence was taken at the time when there was maximal difference between the positive and negative controls (usually 2–5 h) and is expressed as a percentage of the induction observed in the positive control.

### Confocal Microscopy

*C. difficile* strains were grown in 4-well glass chamber slides (BD Falcon, USA) in BHIS + 0.1 M glucose. Chamber slides were incubated at 37°C in anaerobic conditions for 24 hrs – 72 hrs and stained with BacLight live/dead stain mixture (Molecular Probes, Invitrogen), which contains the nucleic acid stains Syto 9 and propidium iodide (staining live bacteria green and dead bacteria red respectively). Following incubation, wells were gently washed twice with PBS 0.1% w/v saponin to remove unattached cells and permeabilise the biofilm. Biofilm samples were incubated with the dye for 15 minutes at 37°C after which the samples were washed twice with PBS. Cells were fixed with 4% paraformaldeyhyde (PFA) for 15 mins. All samples were visualised with a Zeiss Observer LSM 880 confocal scanning microscope at 60x – 100x magnification.

### RNA-seq

Biofilms were grown for 18 h in BHIS + glucose, supernatants were removed and attached biofilms were washed with 1 ml PBS. Biofilms were disrupted and RNA was extracted using Trizol (Invitrogen, UK).5 μg of extracted RNA was treated with RiboZERO^™^ (Illumina, UK) according to the manufacturer’s protocol to deplete rRNA. cDNA libraries were prepared using TruSEQ LT (Illumina, UK) according to the manufacturer’s instructions. Briefly, samples were end-repaired, mono-adenylated, ligated to index/adaptors. Libraries were quantified by bioanalyzer and fluorometer assay. The final cDNA library was prepared to a concentration of 10-12 pM and sequenced using paired end technology using a version-3 150-cycle kit on an Illumina MiSeq^™^ (Illumina, UK).

### RNA-seq Analysis

The paired-end sequencing reads from RNA-seq experiments were mapped against the appropriate reference genome (NC_013316 for *C. difficile* and a de novo assembly using RNA SPAdes v3.9 with default settings [81] from the RNA-sequence reads for *B. fragilis* [Accession number PRJEB29695). The first read was flipped into the correct orientation using seqtk v1.3 (https://github.com/lh3/seqtk) and the reads were mapped against the reference genome using BWA v0.7.5 with the ‘mem’ alignment algorithm [82]. BAM files were manipulated with Samtools v0.1.18 using the ‘view’ and ‘sort’ settings [82]. Sorted BAM and GFF (general feature format) files were inputted into the coverageBed tool v2.27.0 with default settings [83] to gain abundance of each genomic feature. The R package DESeq2 was used with default settings to calculate differential gene expression using a negative binomial distribution model [45]. The data was filtered by applying a cut-off of 1.6 for the fold change and 0.05 for the adjusted *p*-value. All sequencing reads were submitted to the European Bioinformatics Institute (Accession numbers E-MTAB-7486, E-MTAB-7523 and PRJEB29695).

As analysis by BLAST (NCBI) demonstrated species specificity for mapping, coculture samples were mapped to each species reference separately. Initial mapping of the *B. fragilis* strain to a published reference proved unsuccessful, offering a poor rate of alignment of 60%. As the *B. fragilis* strain has not been previously sequenced, and because we were not successful in generating high quality genome sequence, a reference was generated from RNA library of *B. fragilis* using the software rnaSPAdes v3.9 [84] and annotated using Prokka v1.11 (default settings) [85]. The reads from each condition were mapped to their respective reference sequence using BWA v0.7.5 (‘mem’ algorithm) [82, 86] and counted using coverageBed v2.27.0 [83].

Metabolic pathways in *C. difficile* were identified using the KEGG mapper [87], a tool that identifies the function of genes in a published genome. As the *B. fragilis* strain used in this study does not have a published reference genome, blastKOALA [88] was used to search for gene homology within metabolic pathways. Heatmaps were generated from normalised gene expression data outputted from DESeq2, using the online tool Heatmapper [89] using the default settings.

### PCR analysis

16S PCRs were performed using the universal 16S rRNA bacterial primers 27F and 1392R (Table S2). Primers were constructed for prophage genes CDR20219_1208 and CDR20291_1436 (Table S2) to confirm the presence of prophage within *C. difficile* biofilms. PCR was carried out using Fusion High-Fidelity DNA polymerase (NEB, USA) following the manufacturer’s protocol. Samples were heated to 95°C for 5 mins followed by 35 cycles of: 95°C for 30 seconds, 51°C for 30 seconds and 72°C for 30 seconds, after which samples were heated to 72°C for 10 mins.

### eDNA quantification

eDNA was extracted from *C. difficile* biofilms grown in a 24-well plate as described above, using a protocol described in Rice *et al*. [57]. Briefly, the plate was sealed with parafilm and chilled at 4°C for 1 hour. 1 μl 0.5 M EDTA was added to each well and incubated at 4°C for 5 mins. The medium was removed and biofilms were resuspended in 300 μl 50 mM TES buffer (50 mM Tris HCl /10 mM EDTA/500 mM NaCl). The OD_600_ was measured to determine biofilm biomass and the tubes were centrifuged at 4°C at 18,000 g for 5 mins. 100 μl of supernatant was transferred to a tube of chilled TE buffer (10 mM Tris HCl/ 1 mM EDTA) on ice. DNA was extracted using an equal volume of phenol/chloroform/isoamyl alcohol three times. 3 volumes of ice-cold 100% ethanol and 1/10 volumes 3 M sodium acetate were added to the aqueous phase to precipitate the DNA. The DNA pellet was washed with 1 ml ice-cold 70% ethanol, dissolved in 20 μl TE buffer, quantified by Qubit fluorometer (Thermo Fisher).

### Statistical analysis

All experiments were performed in triplicate, with at least three independent experiments performed. Paired student’s t-test was used to determine if differences between two groups were significant, and one way-ANOVA was used to compare multiple groups. Mann-Whitney U tests were used to compare non-parametric data. Fisher’s exact t-test was used to confirm the enrichment of differently regulated genes in prophage regions.

## Supporting information

## Acknowledgments

We thank Gianfranco Donelli, Director, Microbial Biofilm Laboratory, IRCCS Fondazione Santa Lucia, Italy for providing the *Bacteroides fragilis* isolate and Prof Paul Williams, University of Nottingham for providing the *Vibrio harveyi* strain BB170. This was in part supported by a Seed grant from the Wellcome Warwick Quantitative Biomedicine Programme (Institutional Strategic Support Fund: 105627/Z/14/Z).

---

**Figure S1: Analysis of AI-2 production by WT and *LuxS* in planktonic and biofilm growth conditions**.

Strains were grown anaerobically in BHI medium. (**A**) Planktonic aliquots were removed for OD_600_ readings (line) and cell-free supernatants were collected at each time-point (0 – 12 h) and assayed for AI-2 activity using *V. harveyi* BB170 reporter assay (bars). Data shown is the mean of 3 independent experiments in triplicates and error bars indicate SD (**B**) Cell free supernatants were collected from WT and *luxS* biofilms cultured for 24 h and WT planktonic cultures at 8 h. AI-2 activity was measured using *V. harveyi* BB170 reporter assay. Bioluminescence is shown as a percentage of wildtype *V. harveyi* BB120 bioluminescence, which was assumed to be 100 %. Data is representative of 2 independent experiments.

**Figure S2 *C. difficile inhibition in mixed biofilms***

Colony counts from monoculture *C. difficile* and co-culture biofilms with *B. fragilis* after 72 h. Data is representative of 3 independent experiments done in triplicates.

**Figure S3: AI-2 production in mixed biofilms**.

Late log-phase *C. difficile* WT is displayed as a control. Cell-free supernatants were taken from both mono and co-culture biofilms of *C. difficile* (WT and *luxS*) and *B. fragilis*. These were assayed for AI-2 activity using *V. harveyi*. Bioluminescence is shown as a percentage of wild-type *V. harveyi* BB120 bioluminescence, which was assumed to be 100%. Data is representative of 2 independent experiments.

**Figure S4 Cell free *B. fragilis* supernatants do not inhibit *C. difficile***

(**A**) Colony counts were performed at 24 h for WT and WT resuspended in cell-free *B. fragilis* supernatant. (**B**) WT biofilms were grown for 24 h after which the supernatant was replaced with either fresh BHIS + 0.1M glucose, cell-free *B. fragilis* biofilm supernatant or cell-free co-culture biofilm supernatant. Samples were incubated for a further 24 h before colony counts were performed. Data shown is the mean of 3 independent experiments in triplicates and error bars indicate SD, *p < 0.05 as determined by student’s t-test.

**Table S1.**
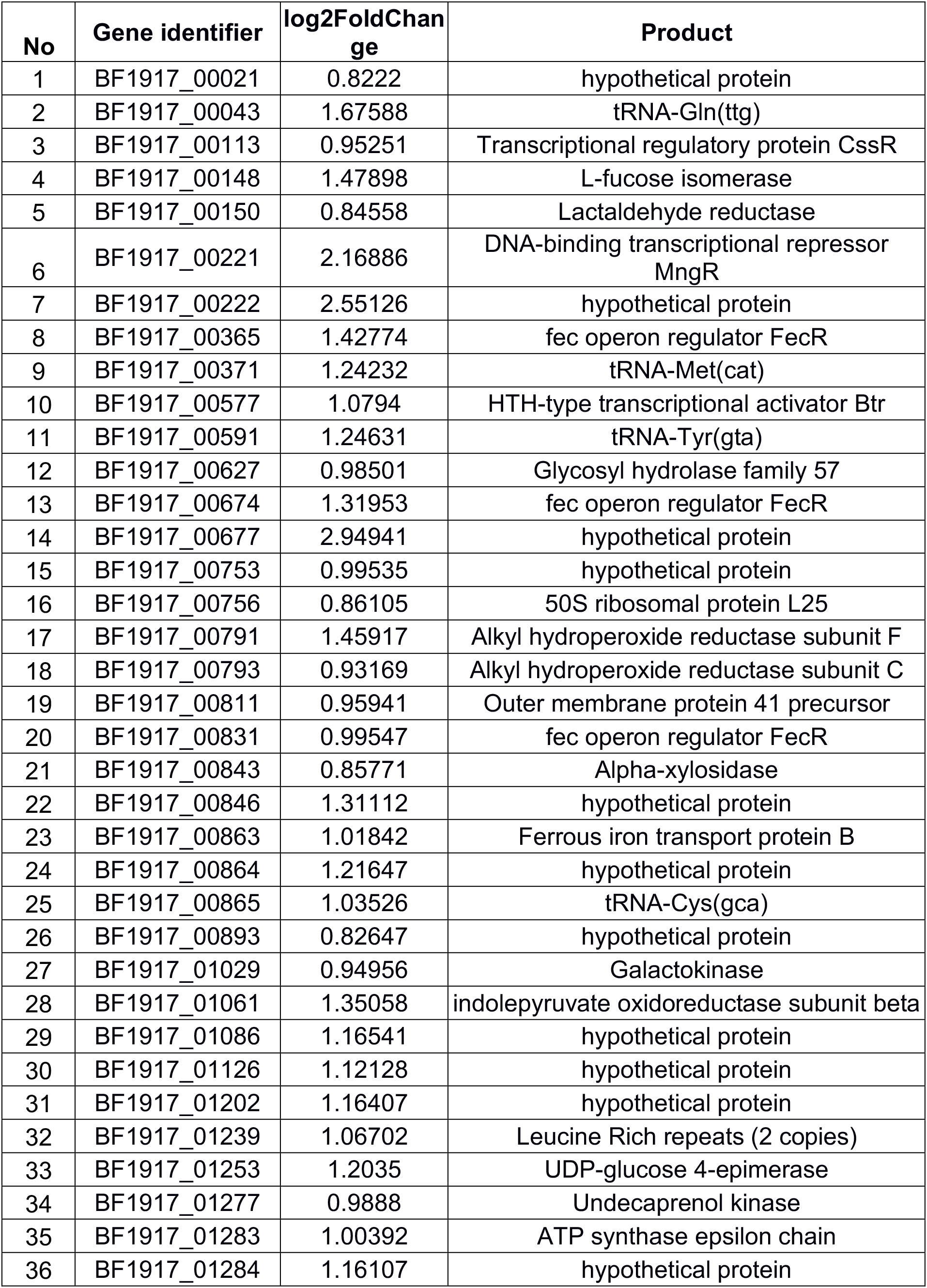
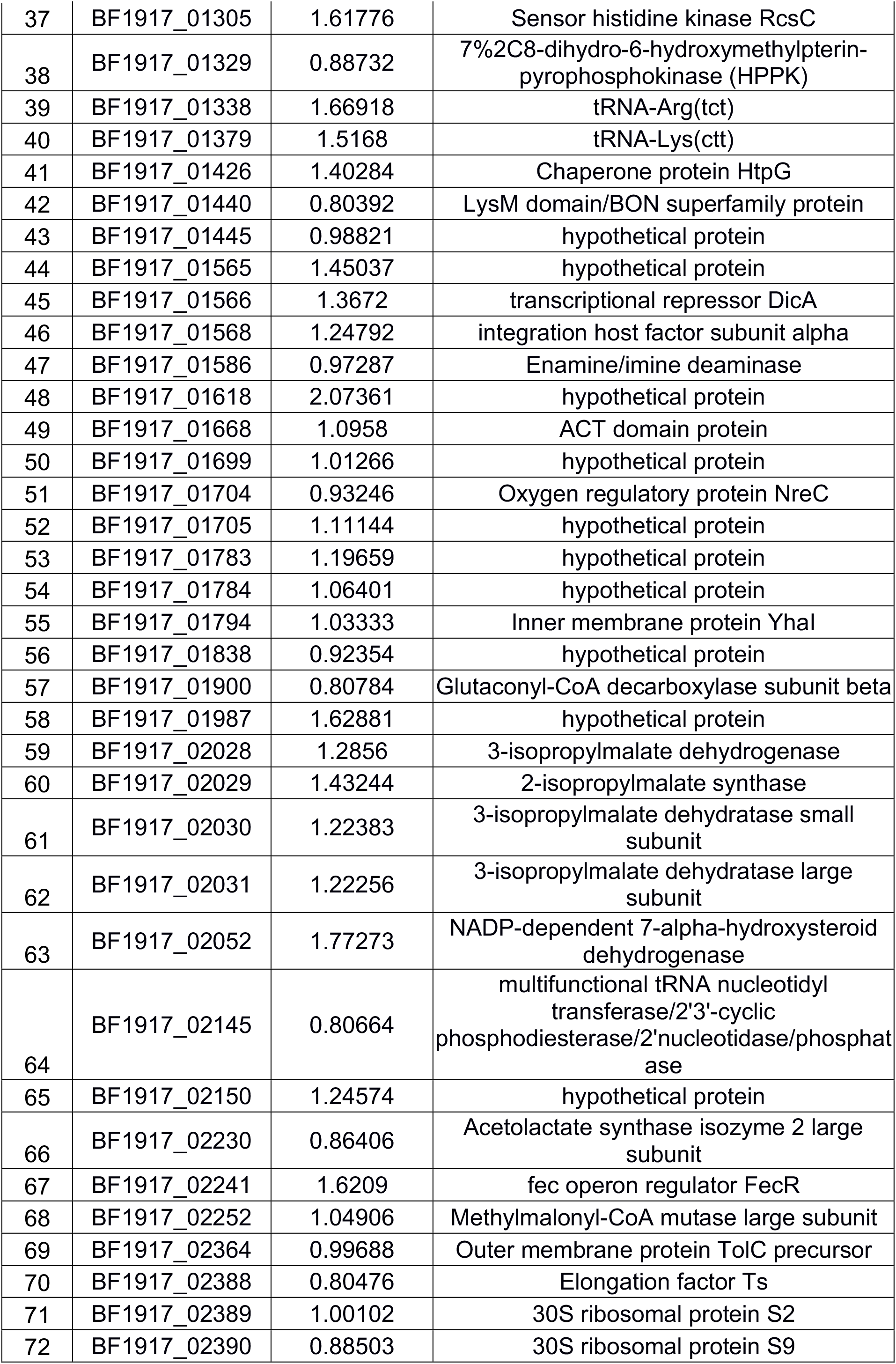
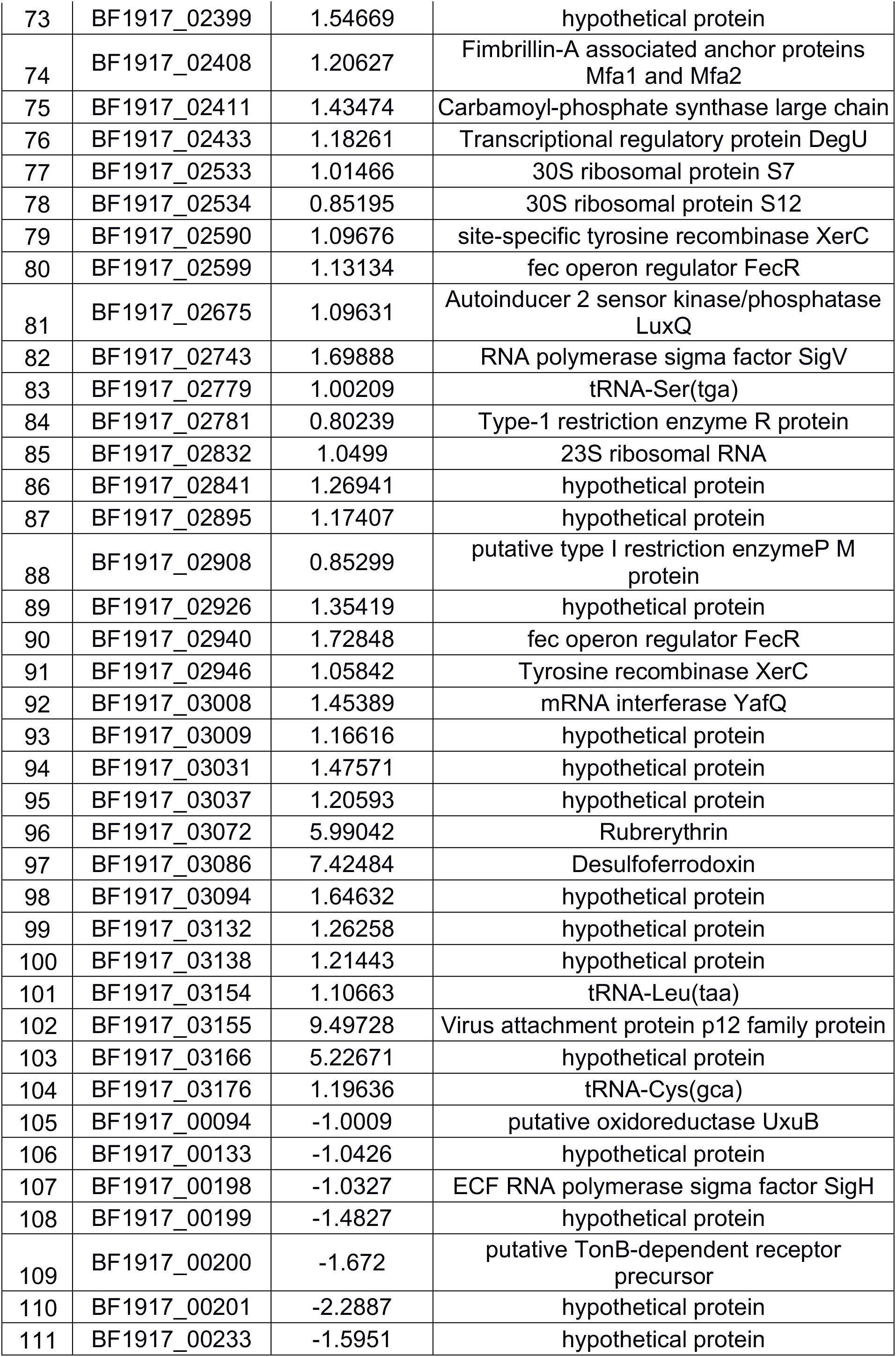
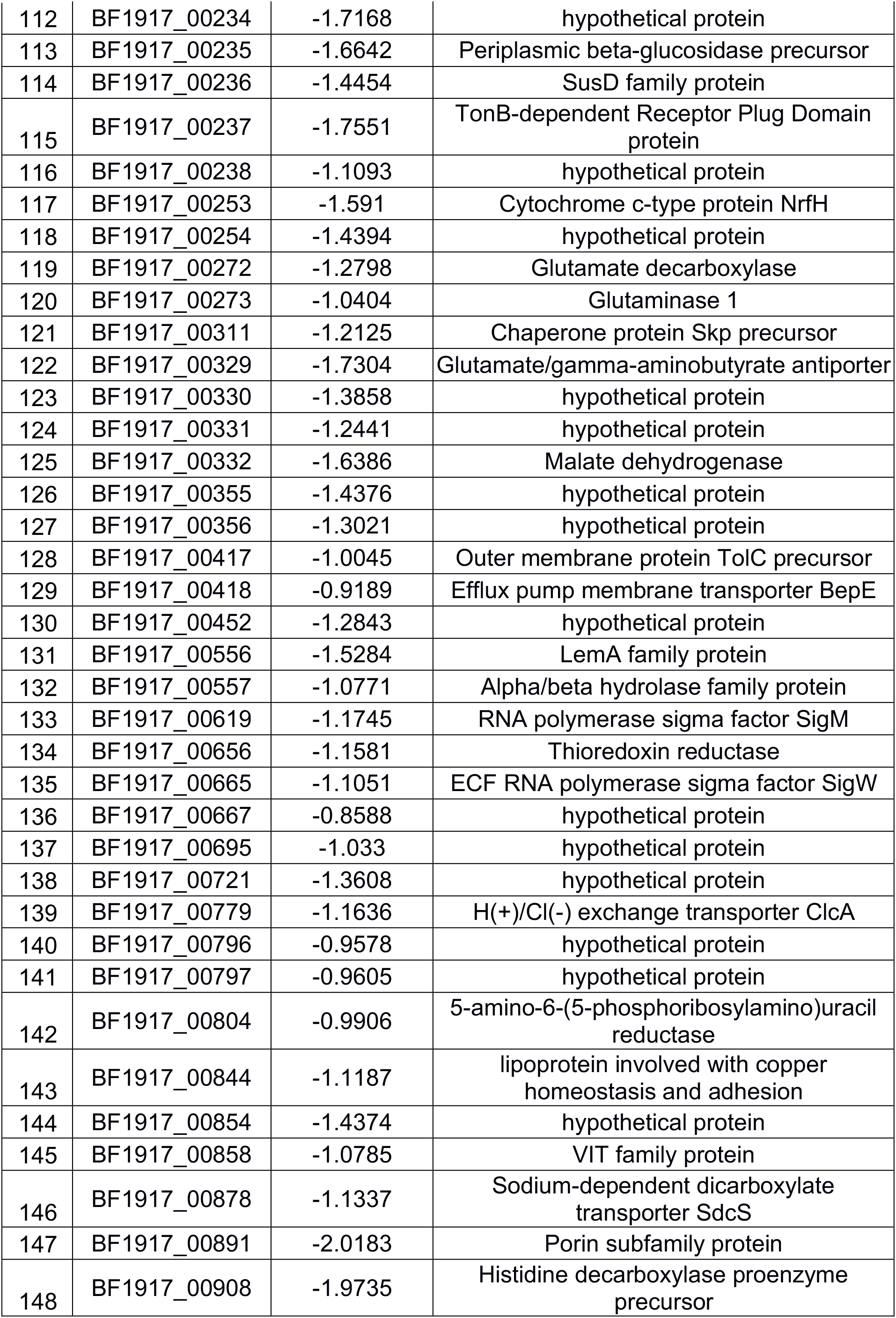
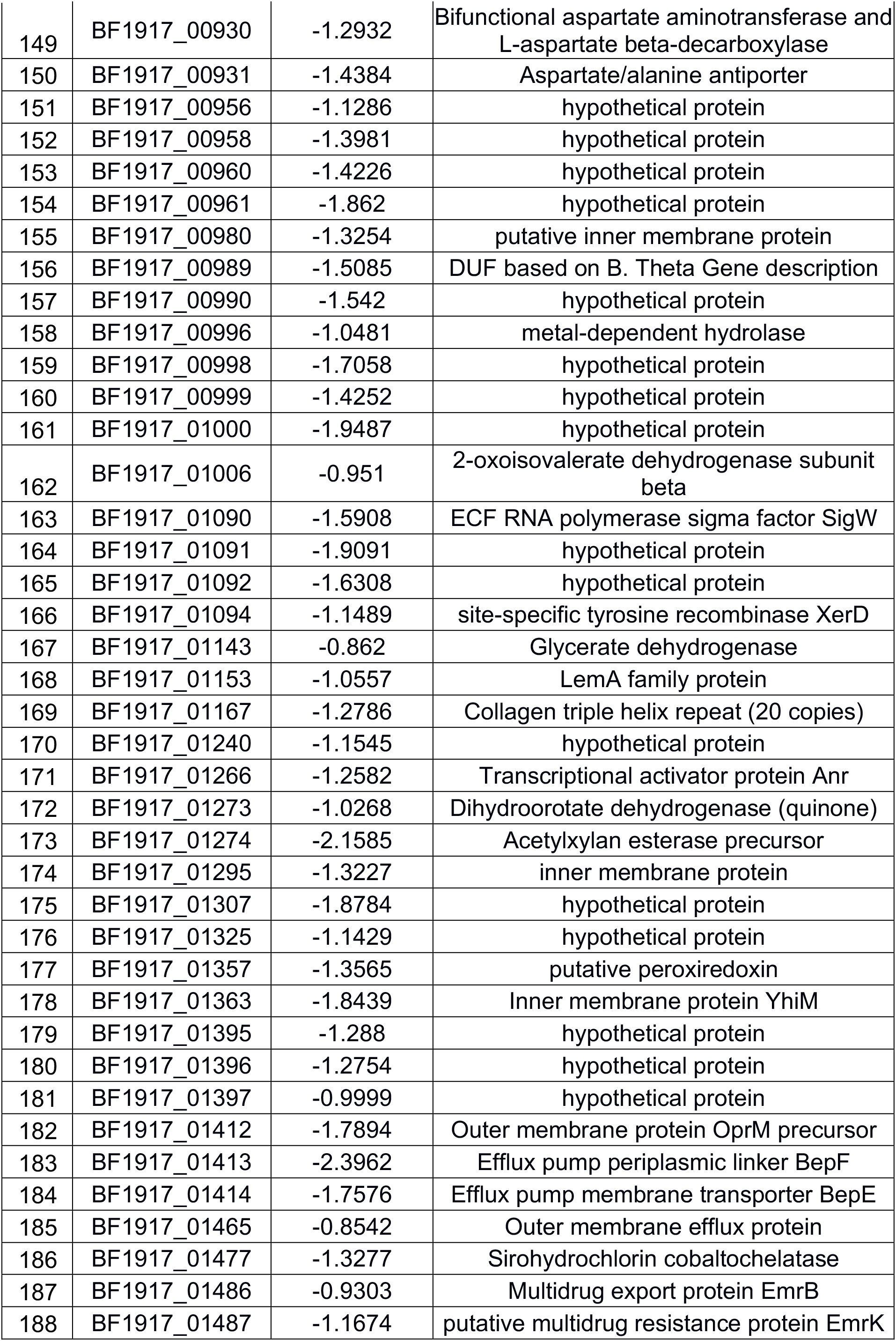
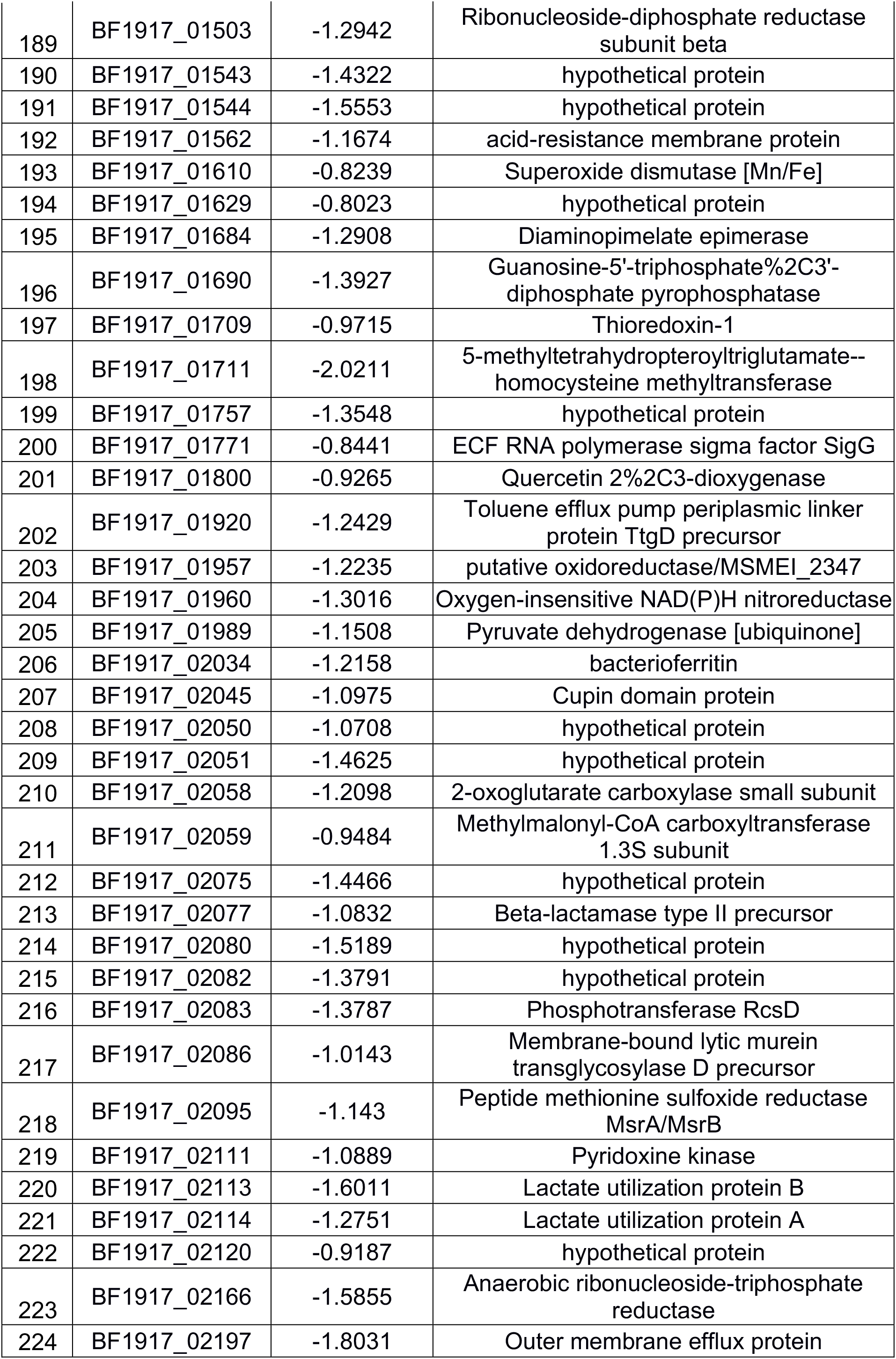
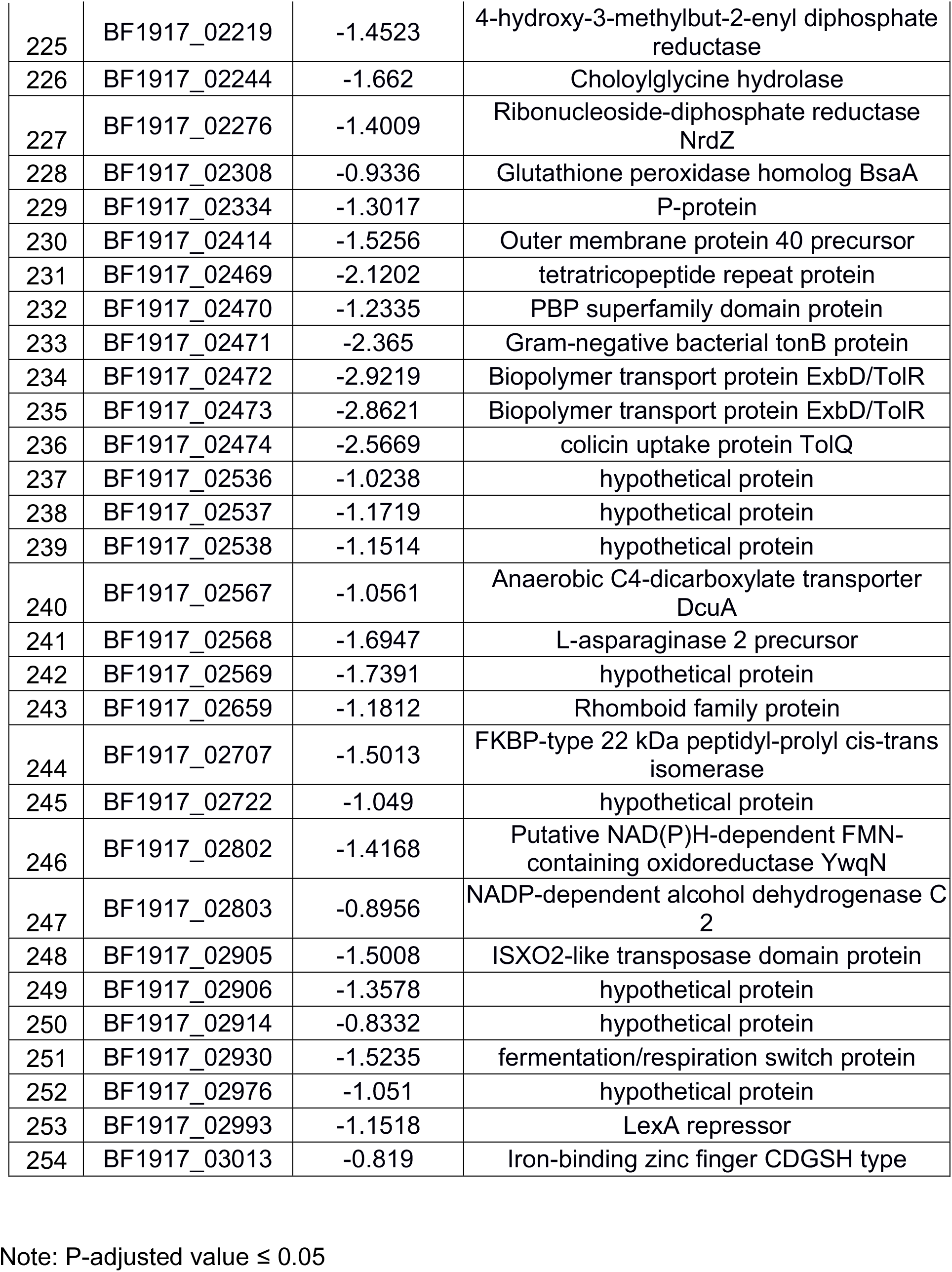
Genes up- and down-regulated in *B. fragilis* co-cultured with both WT and *luxS C. difficile*.

**Table S2.**
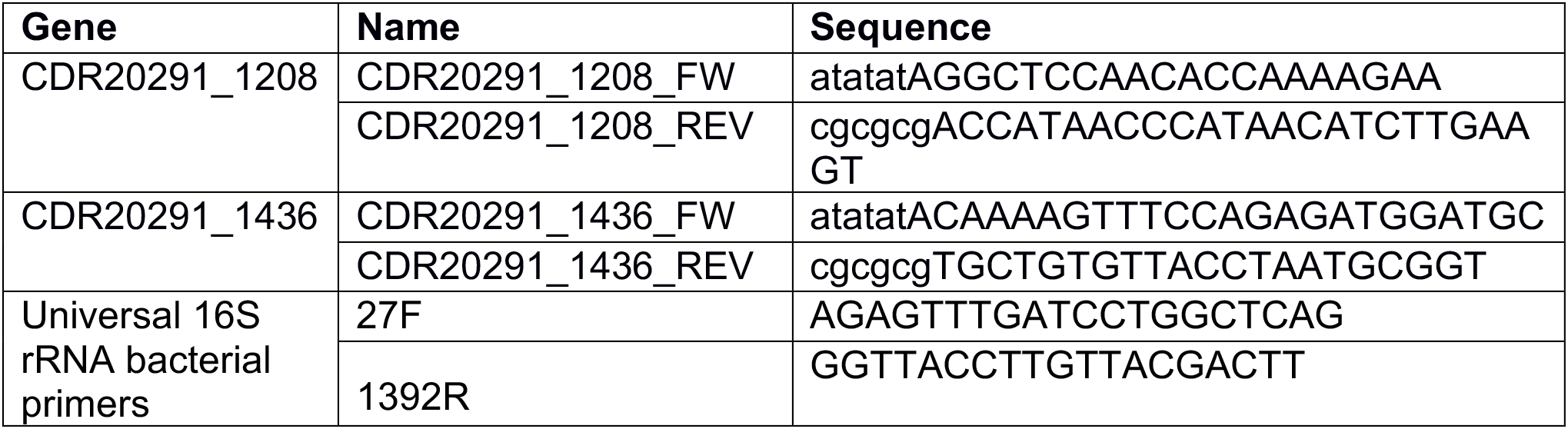
PCR primers for Phage genes identified by RNA-seq

