## Supplementary figures and images for "*Clostridioides difficile* LuxS mediates inter-bacterial interactions within biofilms"

### Supplementary file 1

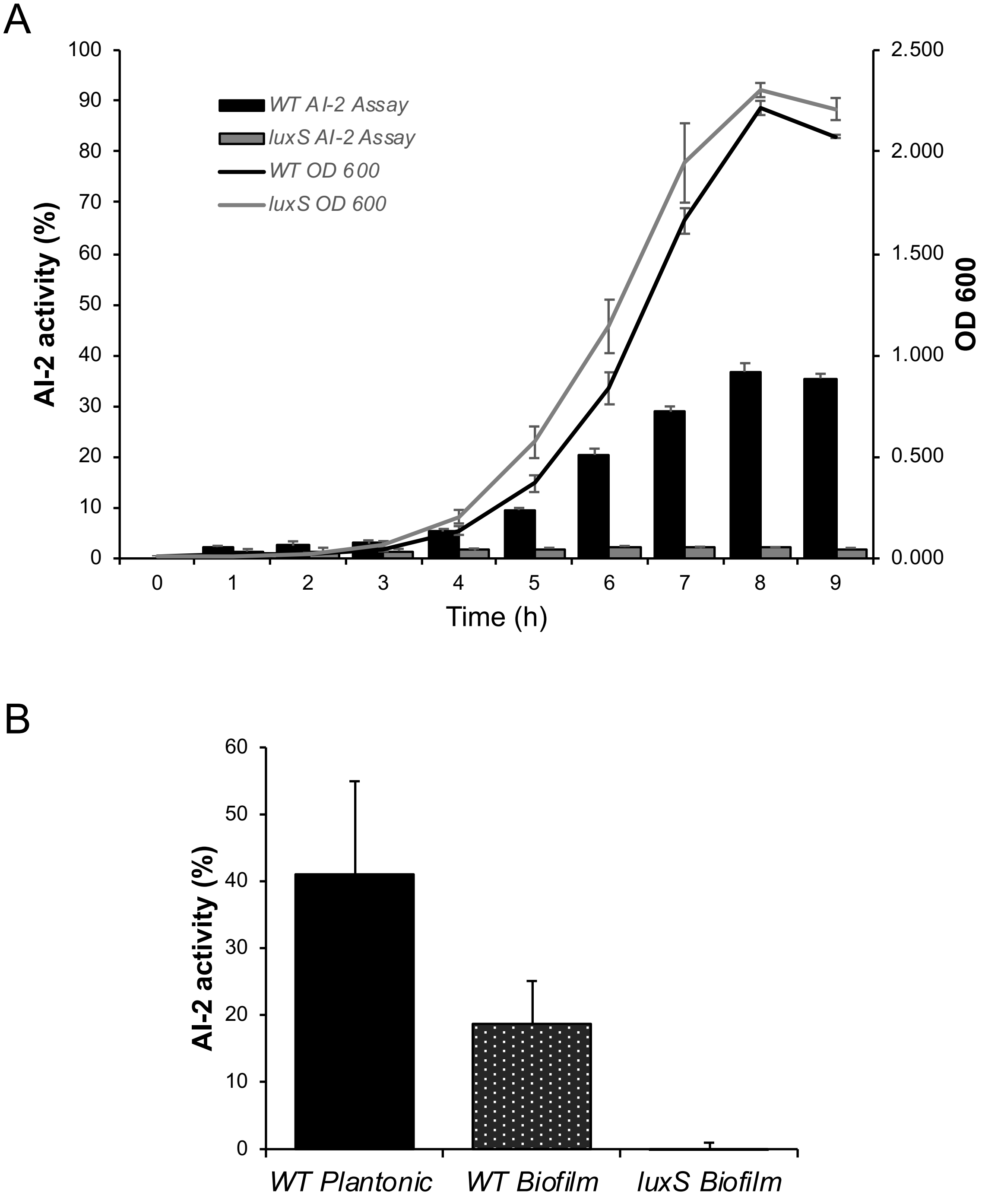

### Supplementary file 2

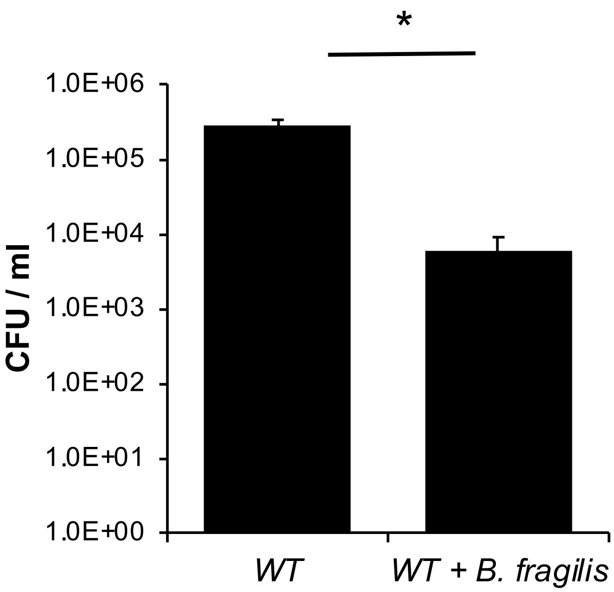

### Supplementary file 3

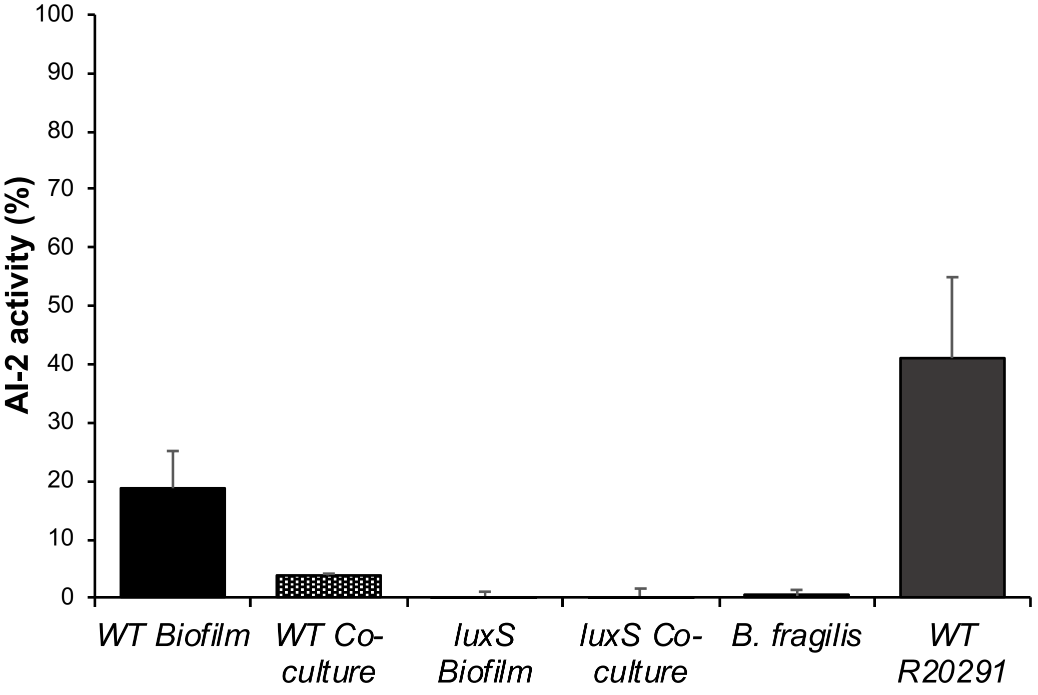

### Supplementary file 4

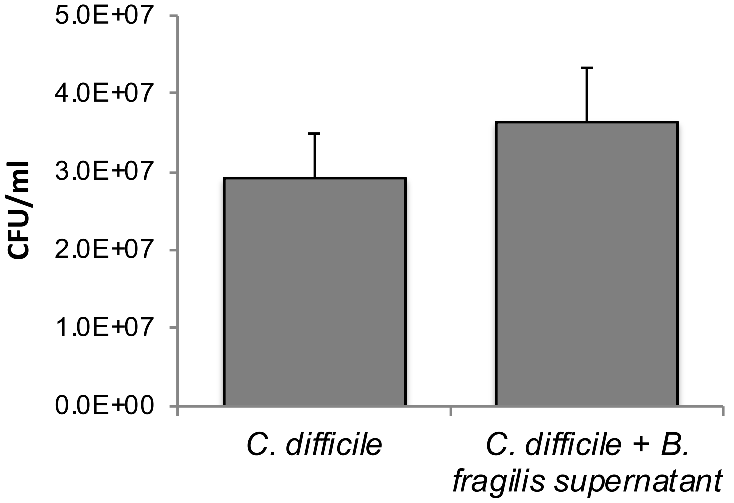

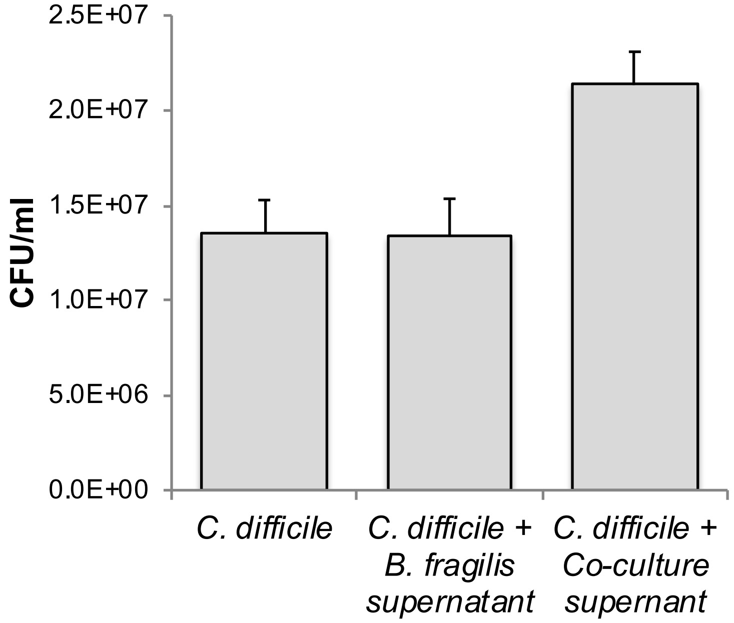
